# Adhesion dynamics regulate cell intercalation behaviour in an active tissue

**DOI:** 10.1101/2021.04.11.439313

**Authors:** Alexander Nestor-Bergmann, Guy B. Blanchard, Nathan Hervieux, Alexander G. Fletcher, Jocelyn Étienne, Bénédicte Sanson

## Abstract

Cell intercalation is a key cell behaviour of morphogenesis and wound healing, where local cell neighbour exchanges can cause dramatic tissue deformations such as body axis extension. Here, we develop a mechanical model to understand active cell intercalation behaviours in the context of an epithelial tissue. Extending existing descriptions, such as vertex models, the junctional actomyosin cortex of every cell is modelled as a continuum morphoelastic rod, explicitly representing cortices facing each other at bicellular junctions. Cells are described directly in terms of the key subcellular constituents that drive dynamics, with localised stresses from the contractile actomyosin cortex and adhesion molecules coupling apposed cortices. This multi-scale apposed-cortex formulation reveals key behaviours that drive tissue dynamics, such as cell-cell shearing and flow of junctional material past cell vertices. We show that cell neighbour exchanges can be driven by purely junctional mechanisms. Active contractility and viscous turnover in a single bicellular junction are sufficient to shrink and remove a junction. Next, the 4-way vertex is resolved and a new, orthogonal junction extends passively. The adhesion timescale defines a frictional viscosity that is an important regulator of these dynamics, modulating tension transmission in the tissue as well as the speeds of junction shrinkage and growth. The model additionally predicts that rosettes, which form when a vertex becomes common to many cells, are likely to occur in active tissues with high adhesive friction.

**SIGNIFICANCE:** Cell intercalation, or neighbour exchange, is a crucial behaviour that can drive tissue deformations, dissipate stress and facilitate wound healing. Substantial experimental work has identified the key molecular players facilitating intercalation, but there remains a lack of consensus and understanding of their physical roles. Existing biophysical models that represent cell-cell contacts with single edges cannot study the continuous dynamics of intercalation, involving shear between coupled cell cortices. Deriving a continuum description of the cell cortex, explicitly coupling neighbouring cortices with adhesions, we define the biophysical conditions required for successful neighbour exchanges. Furthermore, we show how the turnover of adhesion molecules specifies a viscous friction that regulates active tissue dynamics.

## 1 Introduction

In both developing and adult animal tissues, cell rearrangements are a common mechanism by which cells actively drive tissue deformation and passively dissipate stress [1–5]. In epithelia, directed neighbour exchanges between four cells (known as T1 transitions; Figure 1A) are a minimal example of rearrangement that is characterised by the shortening of a shared cell-cell contact, to the point where four cells meet (forming a 4-way vertex), followed by the formation of a new cell-cell contact between previously non-neighbouring cells. This intercalation process can be found throughout development, for example during fish epiboly, mammalian and insect axis extension and hair follicle formation and amphibian and fish neural folding [6–9]. However, surprisingly little is known about the mechanical behaviour of cortical material and adhesions during intercalation. For example, as a junction shrinks, can cortical actomyosin flow around the cell vertex into neighbouring junctions to reduce its length, or must irreversible length changes arise solely from viscous turnover [10, 11]? Furthermore, it is not clear how the properties of adhesion molecules facilitate shearing between coupled cell cortices to allow remodelling, while preserving tissue integrity.

**Figure 1:**
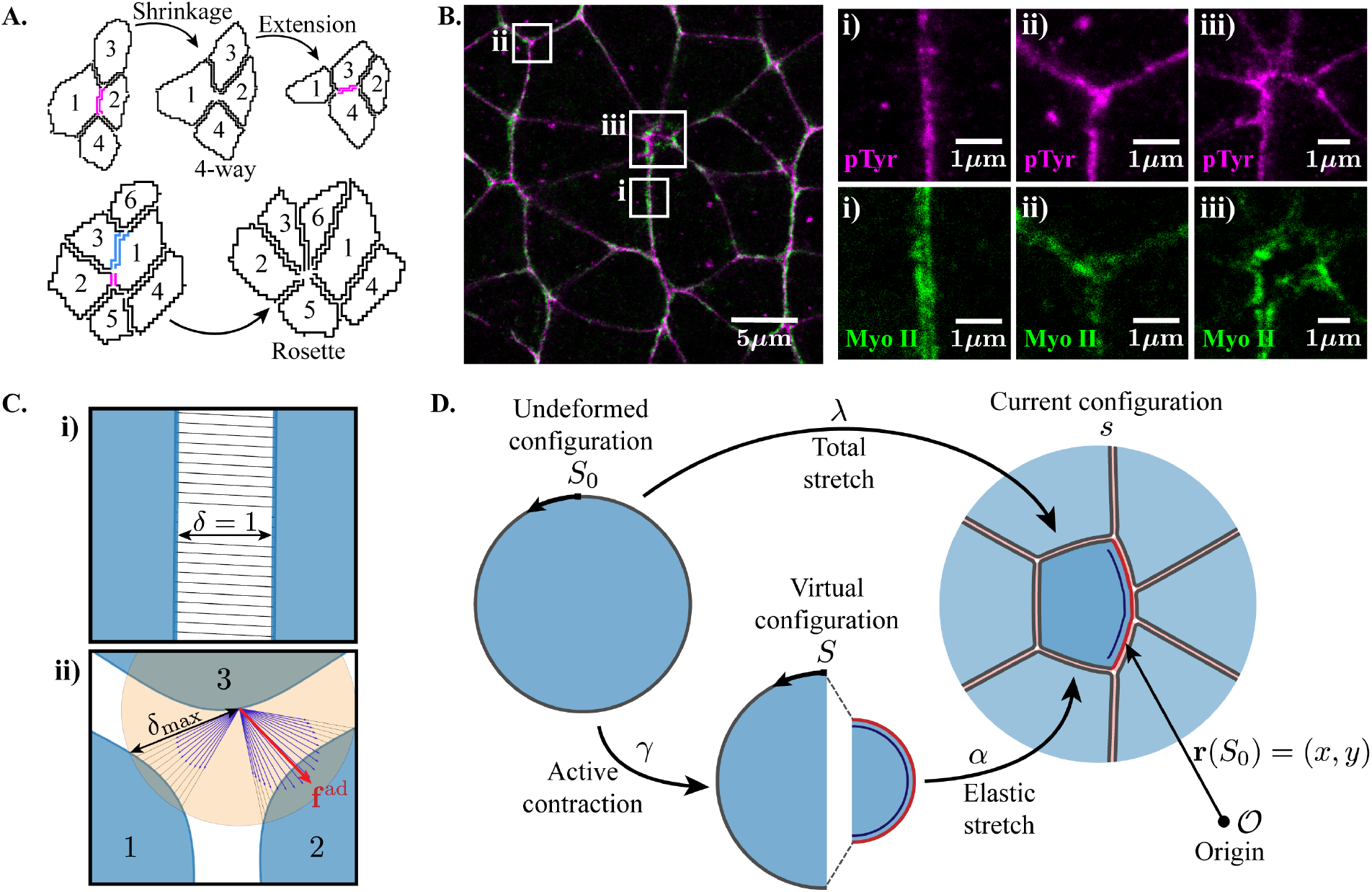
The geometry and mechanics of apposed cortices. **(A)** Example of a T1 transition (top) and a rosette (bottom), segmented from movies of stage 7-8 *Drosophila* embryos [23]. The junctions undergoing shrinkage or elongation are coloured. **(B)** Two-colour STED microscopy image showing the adherens junctions (pTyr; magenta) and cortical myosin II (green) in intercalating cells from stage 8 *Drosophila* embryos. The close ups highlight a bicellular junction (1), a vertex (2) and a rosette center (3). **(C)** Representation of bicellular junctions (i) and vertices (ii) in the apposed-cortex model. Adhesions can bind to neighbouring cortices within *δ*_max_ (orange shading). For the given point on cell 3, blue arrows indicate the inverse lengths of possible adhesions within *δ*_max_. The red arrow represents the mean-field force resulting from the binding kinetics of adhesion when *τ*_ad_ < *τ*_cortex_ (equation (22) in the Supplementary Document). If *τ*_ad_ *≥ τ*_cortex_ unbound cortex nodes bind to the nearest available connection and persist for an average duration *τ*_ad_. **(D)** The three configurations of a cell cortex in the morphoelastic framework: the undeformed configuration (parameterised by *S*_0_) is taken to the virtual/reference configuration (stress-free; parameterised by *S*) by an active contraction, *γ* (the right half of the cortex has *γ <* 1, marked by a dark blue line). The imposed boundary conditions and body loads bring the cortex to the physically realised configuration, called the current configuration (parameterised by *s*), with an elastic stretch, *α*.

Much work has been devoted to identifying and characterising the localisation of the subcellular constituents involved in driving neighbour exchanges across experimental models. Active mechanisms are known to be involved junction shrinkage: contractile myosin II loads the junctional cortex in invertebrate [12–17] and vertebrate models [18–20]. However, the physical roles that these subcellular features take and whether they are strictly necessary, or merely supportive, are not known. It is currently also unclear whether such active mechanisms are required to drive the growth of a new junction, or if extension is energetically favourable and follows passively. Following shrinkage, before extension can occur, the 4-way vertex must be resolved. Delays or failures in resolution can lead to the formation of higher-order vertices, shared by many cells, known as rosettes (Figure 1A) [21, 22]. A higher prevalence of rosette structures has been linked to perturbations in the mechanical properties of tissues and defects in tissue reshaping during morphogenesis [23, 24]. However we have little information about what defines the 4-way resolution timescale and whether it is actively tuned to prevent topological defects in a tissue.

Discrete vertex-based models are perhaps the most popular models of epithelial tissues [2, 25–28]. These models are simple and computationally inexpensive, yet successfully capture a remarkable variety of tissue behaviour. However, neighbour exchanges must be implemented manually in a discrete framework [26], if they are explicitly represented [29]. This can lead to unwanted flipping between junction shrinkage and growth [30]. Discrete models can successfully explore the biochemical conditions that can drive junction shrinkage [15, 31]. However, since intercalation is fundamentally a continuous process, they cannot explore how 4-way vertices are resolved or the mechanisms regulating their timescales.

With few exceptions [32], vertex-based models most often use a single edge to represent a bicellular junction [33, 34], without explicit representation of the apposed actin cytoskeletons and adhesion bonds coupling them. This implicitly imposes that neighbouring cortices cannot move apart or shear relative to one-another – a crucial aspect of intercalation. The separated cortices of cells in the *Nematostella vectensis* have been modelled using a discrete network of multiple springs [35]. However the discrete model is limited to the behaviour of springs in series. The immersed boundary method successfully models apposed cortices as continuum structures [36], providing insight to tumour growth and limb bud morphogenesis [37, 38]. However, the model requires the presence of an unphysical incompressible fluid between cell cortices, requiring artificial source and sink terms to control growth [39]. A fluid-free cell boundary model, that is more faithful to existing physical quantities, is likely to be more appropriate for simulating epithelial tissue behaviour.

In this paper, we present an apposed-cortex model of an epithelial tissue that has direct access to subcellular mechanical constituents. The junctional cortex is modelled as a continuum morphoelastic rod with explicit adhesions between the cortices of different cells. Shear between apposed cortices and the displacement of vertices along the cortices are emergent features of the model. The model is used to understand the minimal conditions under which cell neighbour exchanges (junction shrinkage, resolution and extension) can be driven by active subcellular contractility in the junctional actin network. We demonstrate that adhesion dynamics are a key feature in regulating the dynamics of an active tissue.

## 2 Results

### 2.1 Introduction to the apposed-cortex model: passive mechanics

Across multiple tissue types, actomyosin and adhesion molecules, such as E-cadherin, are key mechanical constituents that modulate tissue dynamics [4, 7]. In order to gain a mechanistic understanding of how their properties regulate neighbour exchanges, a model must have explicit access to these components. This requires consideration of the geometric and topological configurations in which cells, and thereby actomyosin and adhesion molecules, are organised. We confirm, using high-resolution microscopy (STED) in the *Drosophila* germ-band, that myosin loads each apposed cortex in a bicellular junction separately (Figure 1Bi). Furthermore, direct access to adhesion dynamics requires modelling each cell cortex explicitly with adhesion bonds coupling the cortices of neighbouring cells (Figure 1C).

We begin by considering the passive mechanical properties of a cell and assume that the cortex is the leading mechanical driver of cell behaviour [40]. Each cell cortex is modelled as an extensible, unshearable and torsion-free rod [41], whose centreline is parameterised by a Lagrangian coordinate, *S*, in the reference (stress free) configuration (Figure 1D). The cortex is assumed to have resistance to bending and extension, with dimensionless elastic mechanical energy, 𝒰, given by

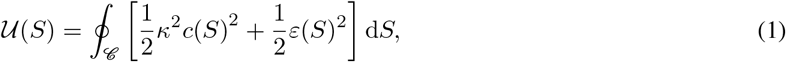

where *ε* = *α −* 1 is the strain with elastic stretch, *α*, and *c/α* is the Frenet curvature; 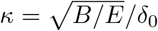 is a dimensionless mechanical parameter representing the ratio of bending, *B*, to extensional, *E*, modulus, relative to a reference length, *δ*_0_. At rest, an isolated segment of cortex would lie as a straight rod.

We allow the elastic forces generated within the cortex to be dissipated viscously. Cortex relaxation is mainly driven by actin turnover time, on timescales of *∼* 50 s [10, 42]. Active cell rearrangement occurs on tens-of-minutes timescales [17]. Given that the cortex relaxation time is significantly shorter than the timescale over which rearrangement occurs, we assume that the cortex behaves as a purely fluid material and update the rest length to the current length, between mechanical equilibria, at every simulation step.

In addition to the passive constitutive material properties of the cortex, we consider forces acting on the cortex due to adhesions. Adhesion is modelled as a single agent that accounts for the composite effect of all molecules associated with the adhesion complex, such as E-cadherin, *α*- and *β*-catenin and vinculin [43]. Adhesion bonds acting over a unit length of cortex, coupling it to another cortex, are each modelled as a simple spring, with energy (Figure S1A)

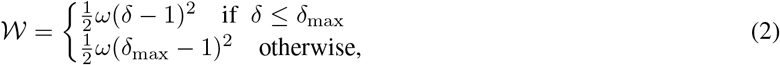

where *ω* is the stiffness of an adhesion bond, times the lineic density of bonds, and scaled by the extensional modulus, *E*, of the cortex; *δ* is the dimensionless length of the adhesion bond (the distance between the two cortices that it is attached to; Figure 1C). Lengths have been scaled relative to the rest length of an adhesion bond, *δ*_0_. Adhesion bonds break when *δ > δ*_max_. However, given that adhesion recovery times are seen to be very fast (*∼* 20 s [44, 45]), we impose that all cortex locations without adhesion bonds immediately attempt to re-bind to neighbouring cortices lying within distance *δ*_max_. From Figure 1C, we see that a vertex in this model is the geometric point where three or more cells are coupled by adhesions, rather than a material point.

Adhesion bonds are additionally given a timescale representing their average lifetime, *τ*_ad_. This timescale corresponds to how many simulation steps a bond persists for, on average, before it disassociates. The cortical timescale, *τ*_cortex_, is defined by viscous actin turnover at every simulation step. When *τ*_ad_ < *τ*_cortex_ adhesion forces work in the mean-field, taking the average force of all possible binding pairs within *δ*_max_ (detailed in the Supplementary Document; Figure 1C, vertex, shows an example mean-field force). When *τ*_ad_ *≥ τ*_cortex_ adhesion bonds associated to specific cortex locations persist for multiple simulation steps, with an average fraction 1 − 1*/τ*_ad_ turning over every time step. The effect of *τ*_ad_ on tissue dynamics is explored in later sections.

Accounting for cortical material forces with adhesions, the balance of linear momentum in the reference configuration requires

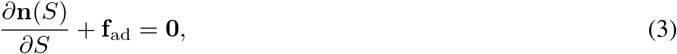

where **f**_ad_ is the external force due to adhesions. The cortical force, **n**, is given by (see the Supplementary Document)

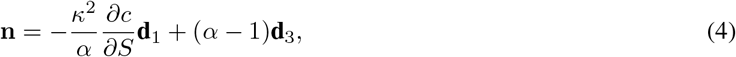

where (**d**_1_, **d**_3_) are orthonormal vectors in the normal and tangential directions along the cortex (Figure S1B). We can generate a tissue with multiple cell cortices, coupled by adhesions, each satisfying (3).

### 2.2 Vertex geometry is determined by the cortex bending stiffness relative to adhesion strength

The constitutive properties of the cortex are determined by the bending, *B*, and extensional, *E*, moduli, captured by their dimensionless ratio, *κ*^2^. This ratio relates to the length of the curved opening around cell vertices, which can be used to parameterise the model. Consider a single cell enclosed by a hexagonal boundary (Figure 2A). The size of the opening at the vertices of the hexagonal boundary is determined by *ω* and *κ*: increasing *κ* increases the penalty for having sharp corners (large curvature), requiring stronger adhesion, *ω*, to close the vertex. The size, *δ*^vert^, of the opening at vertices is therefore a geometric feature that characterises the passive mechanical properties of a tissue. Mapping this geometric feature across (*κ, ω*) parameter space we identify isolines of constant *κ*^2^*/ω*, with equal *δ*^vert^ (Figure 2B). The cortex extensional modulus, *E*, cancels out in the ratio *κ*^2^*/ω*, such that the geometry around vertices is given by the ratio of cortex bending modulus to adhesion modulus.

**Figure 2:**
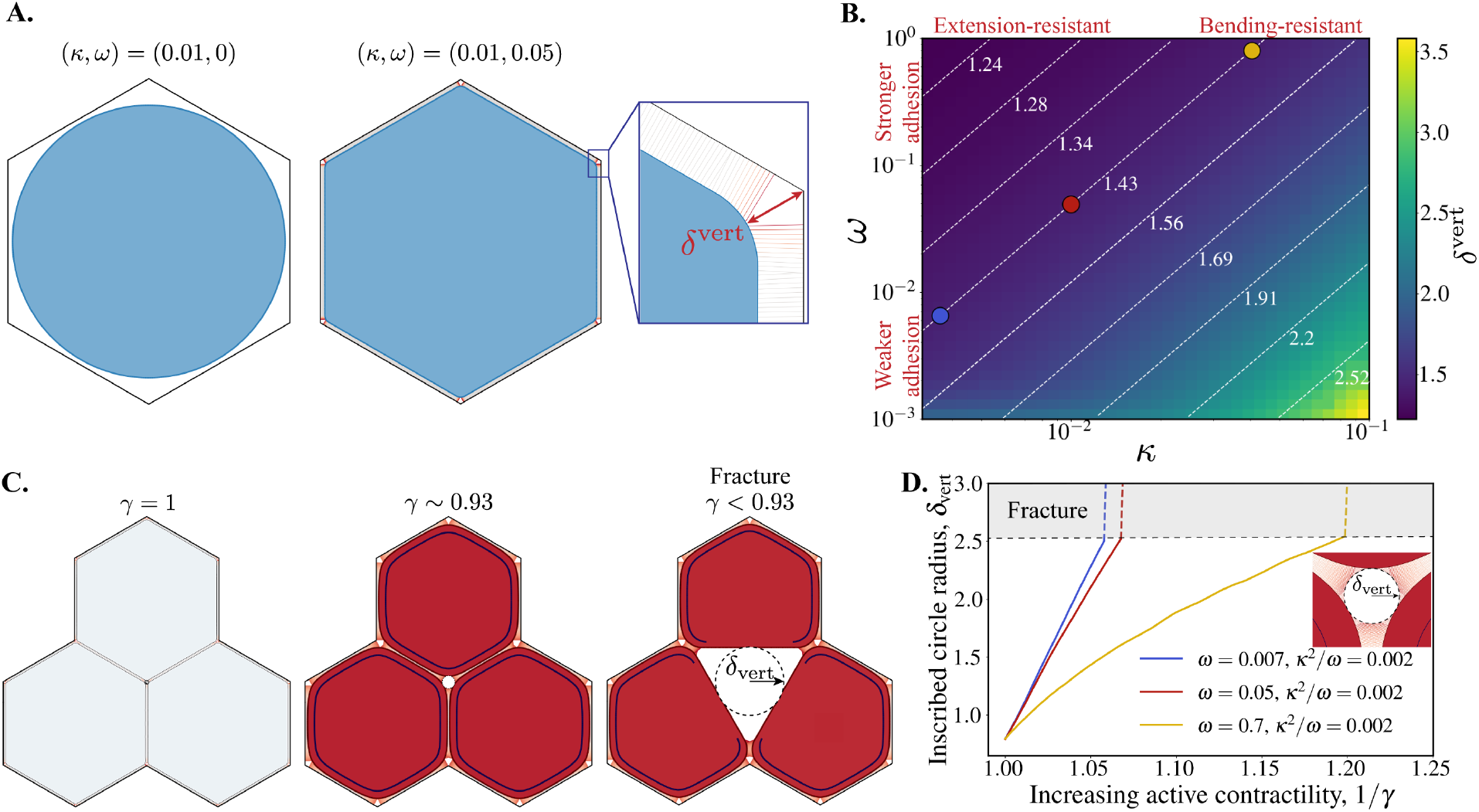
Passive and Active mechanics in the apposed-cortex model. **(A)** A circular cell cortex is initialised with no adhesions (*ω* = 0). The adhesion strength can be quasi-statically increased to pull the cell towards the hexagonal boundary and close the vertices. **(B)** Map of parameter space showing the opening at a vertex, *δ*^vert^, (see **A**) relative to the parameters *κ* and *ω* on a log scale. For each (*κ, ω*), the cell was recurrently relaxed to equilibrium with their rest lengths updated to the current length at every step, repeated until the cortex length change was less than 1 × 10^−4^. Dashed white lines show isolines of fixed *δ*^vert^, corresponding to constant *κ*^2^*/ω*. Coloured circles show the parameter samples along the isoline used in **D. (C)** Snapshots from a 3-cell simulation with (*κ, ω*) = (0.01, 0.05) and increasing active cortical contractility in all cells (*γ* decreasing from 1; see also Movie 1A). The tissue fractures when *γ <* 0.93. The dashed line shows a circle inscribed within the vertex, with radius *δ*_vert_. **(D)** Radius of the vertex-inscribed circle vs. the inverse magnitude of active contractility for three parameter samples on the *κ*^2^*/ω* = 0.002 isoline in passive parameter space, where *δ*_vert_ is constant (coloured dots in **B**).

To our knowledge, there are no explicit measurements of the size of vertex openings in the literature. Our STED imaging of adherens junction in the germ-band does not show any significant opening at vertices (Figure 1B, panel ii, pTyr). Higher resolution imaging with electron microscopy of other fly tissues also suggests that vertices are tightly closed (see e.g. Figure 7 of [46]). We therefore choose order-of-magnitude estimates *κ* = 0.01, *ω* = 0.05, which give *δ*^vert^ *∼* 1.43 (relative to the bicellular spacing; Figure 2B, red marker). Figure S2A shows some representative vertex structures across *κ*^2^*/ω* samples.

### 2.3 Active mechanics in the apposed-cortex model

We use the theory of morphoelasticity to incorporate generalised active stresses in the cell cortex. We must consider three configurations for the cortex: first, the initial configuration in which the cortex is undeformed and has no active stress; second, the reference configuration in which the cortex is unstressed, though subjected to active (pre)stress that deforms it. This is a virtual configuration that cannot, in general, be achieved in Euclidian space; third, the current configuration, which is the physically realised configuration that the cortex adopts in response to active stresses and external forces. These configurations are visualised in Figure 1D. The cortex is taken from its undeformed configuration, parameterised by *S*_0_, to the virtual configuration, parameterised by *S*, by an active stretch:

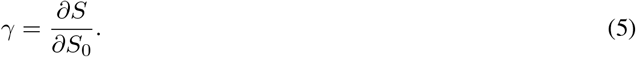

The active stretch represents local changes in material length, with *γ >* 1 for active growth and *γ <* 1 representing active contraction. Though we use traditional terminology, referring to *γ* as a stretch, we deal only with active contraction in a tissue. Setting *γ <* 1 could represent loading regions of the cortex with active myosin, the main driver of contractility in many tissues. The effect of loading contractile machinery on the cortex can hence be conceptualised as substituting cortex segments with segments that have a smaller rest length. The elastic stretch is then the local change of length between the virtual configuration and the current configuration, *s*, due to external physical constraints:

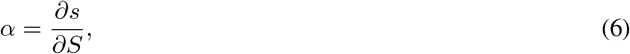

from which elastic stresses arise. We next perform simulations with active contractility. The parameters used for each simulation are given in Table 1.

**Table 1:** Table showing parameters used in simulations. ^†^ For this simulation with asymmetric contractility across some bicellular junctions, cells 1 & 2 were given *γ* = 1 − 0.01 and cells 3 & 4 were set with *γ* = 1 − 0.0025.

| Simulation | $\kappa$ | $\omega$ | $\tau_{\text{ad}}$ | $\gamma$ | $\gamma_0$ | Prestress cell pairs | $\delta_{\text{max}}$ | $\delta_\gamma$ |
| --- | --- | --- | --- | --- | --- | --- | --- | --- |
| Movie 1A | 0.01 | 0.05 | $\tau_{\text{cortex}}$ | $1 \rightarrow 0.9$ | 1 | $\{1, 2\}, \{1, 3\}, \{2, 3\}$ | 4 | 4 |
| Movie 1B | 0.01 | 0.05 | $\tau_{\text{cortex}}$ | $1 - 0.005$ | 1 | $\{1, 2\}$ | 4 | 4 |
| Movie 2A | 0.01 | 0.05 | $10 \tau_{\text{cortex}}$ | $1 - 0.04$ | $1 - 2 \times 10^{-4}$ | $\{1, 2\}$ | 4 | 4 |
| Movie 2B | 0.01 | 0.05 | ∞ | $1 - 0.04$ | $1 - 2 \times 10^{-4}$ | $\{1, 2\}$ | 4 | 4 |
| Movie 3A | 0.01 | 0.05 | $< \tau_{\text{cortex}}$ | $1 - 0.04$ | $1 - 2 \times 10^{-4}$ | $\{1, 2\}$ | 4 | 4 |
| Movie 3B | 0.01 | 0.05 | $< \tau_{\text{cortex}}$ | $1 - 0.04$ | $1 - 2 \times 10^{-4}$ | $\{1, 2\}$ | 4 | 4 |
| Movie 3C | 0.01 | 0.05 | $10 \tau_{\text{cortex}}$ | $1 - 0.04$ | $1 - 2 \times 10^{-4}$ | $\{1, 2\}$ | 4 | 4 |
| Movie 3D | 0.01 | 0.05 | $< \tau_{\text{cortex}}$ | $1 - 0.04$ | $1 - 2 \times 10^{-4}$ | $\{1, 2\}$ | 4 | 4 |
| Movie 3E | 0.01 | 0.05 | $10 \tau_{\text{cortex}}$ | $1 - 0.04$ | $1 - 2 \times 10^{-4}$ | $\{1, 2\}$ | 4 | 4 |
| Movie 4A | 0.01 | 0.05 | $10 \tau_{\text{cortex}}$ | $1 - 0.01$ & $1 - 0.0025^\dagger$ | $1 - 2 \times 10^{-4}$ | $\{1, 2\}, \{1, 3\}, \{1, 4\}, \{2, 3\}, \{2, 4\}$ | 4 | 2 |
| Movie 5A | 0.01 | 0.05 | $\tau_{\text{cortex}}$ | $1 - 0.04$ | $1 - 2 \times 10^{-4}$ | $\{1, 2\}, \{1, 3\}$ | 4 | 4 |
| Movie 5B | 0.01 | 0.05 | $100 \tau_{\text{cortex}}$ | $1 - 0.04$ | $1 - 2 \times 10^{-4}$ | $\{1, 2\}, \{1, 3\}$ | 4 | 4 |

### 2.4 Active contractility affects vertex stability and can drive material flow between junctions

The geometry of cell vertices, set by *κ*^2^*/ω*, is perturbed when cells are mechanically active. Simulating isotropic whole-cortex contractility (setting *γ <* 1) in a minimal 3-cell tissue, we find that active stresses perturb vertex stability. For sufficiently strong active contractility, the vertex opening is forced beyond the maximum adhesion binding length, *δ*_max_, and the tissue fractures (Figure 2C and Movie 1A). Tracking the size of the vertex opening as active contractility is increased, we see that larger values of the adhesion strength, *ω*, produce more stable vertices that can sustain more active stress before fracturing (Figure 2D). Cell vertices are thus potential sites of weakness in a tissue and there is a critical contractility threshold, beyond which the tissue will fracture.

If the forces acting at a vertex are not isotropic, in addition to displacing the vertex, cortical material can flow between cell junctions, across the vertex (Movie 1B and Figure S2B). Indeed, from the view point of a given cell, a vertex can slide along the cortex. Adhesions disconnect along the junction that is receding, while new adhesions form with the cortex of the cell that is advancing. A pair of neighbouring bicellular junctions can thus commensurately increase and decrease their lengths by passing material between each other, across the vertex (Figure S2C). Importantly, this behaviour cannot be manifested by models where junctions behave as spring-dashpot elements connected at vertices (Figure S3A). There, shrinkage can happen from contraction and viscous dissipation only. In this model, the cell cortex behaves as an extensible, viscous rope that is anchored by cell-cell adhesions. We can conceptualise this as a rope surrounding pulleys at vertices, with a friction that resists material flow between junctions (Figure S3B). Now, shrinkage can be achieved by both contraction and material flow between junctions. In the model presented here, the friction associated with this shear flow depends on the adhesion timescale and is studied in further detail below.

### 2.5 Active contractility can drive complete junction removal

Active neighbour exchanges are often driven by the enrichment of subcellular components on a subset of bicellular junctions [4]. In particular, contractile myosin II motors are commonly thought to drive junction shrinkage. We simulate this process by localising active stress on a single bicellular junction in a minimally sized tissue (Figure 3). The choice of fourteen cells allows all cells connected to the shrinking junction to be surrounded by other cells, whilst keeping the computational cost to a minimum. In order to reproduce the overall tissue tension revealed by laser ablations [47, 48], we impose a low level of prestress, setting *γ* = *γ*_0_ = 1 *— ϵ* for *ϵ ≪* 1, in all cell cortices. Recent experimental evidence suggests that cells in the *Drosophila* germ-band enrich cortical myosin using a neighbour ‘identity’-sensing mechanism [17, 49–51]. In particular, cells have been found to alter the material properties of their cortex in response to genetically-specified asymmetric localisation of cell surface receptors between the apposed cortices of a bicellular junction. We mimic this mechanism by making the local level of active contractility in a cortex dependent on the identity of the cells with whom it can form such a coupling. In a similar manner to adhesion, the identity sensing couplings have a maximum binding range *δ*_*γ*_. Unless otherwise stated, we set *δ*_*γ*_ = *δ*_max_ for simplicity. In this case, we simulate the active contraction of the bicellular junction between cells 1 and 2 by setting *γ <* 1 in the portions of their cortices where they can share adhesion bonds.

**Figure 3:**
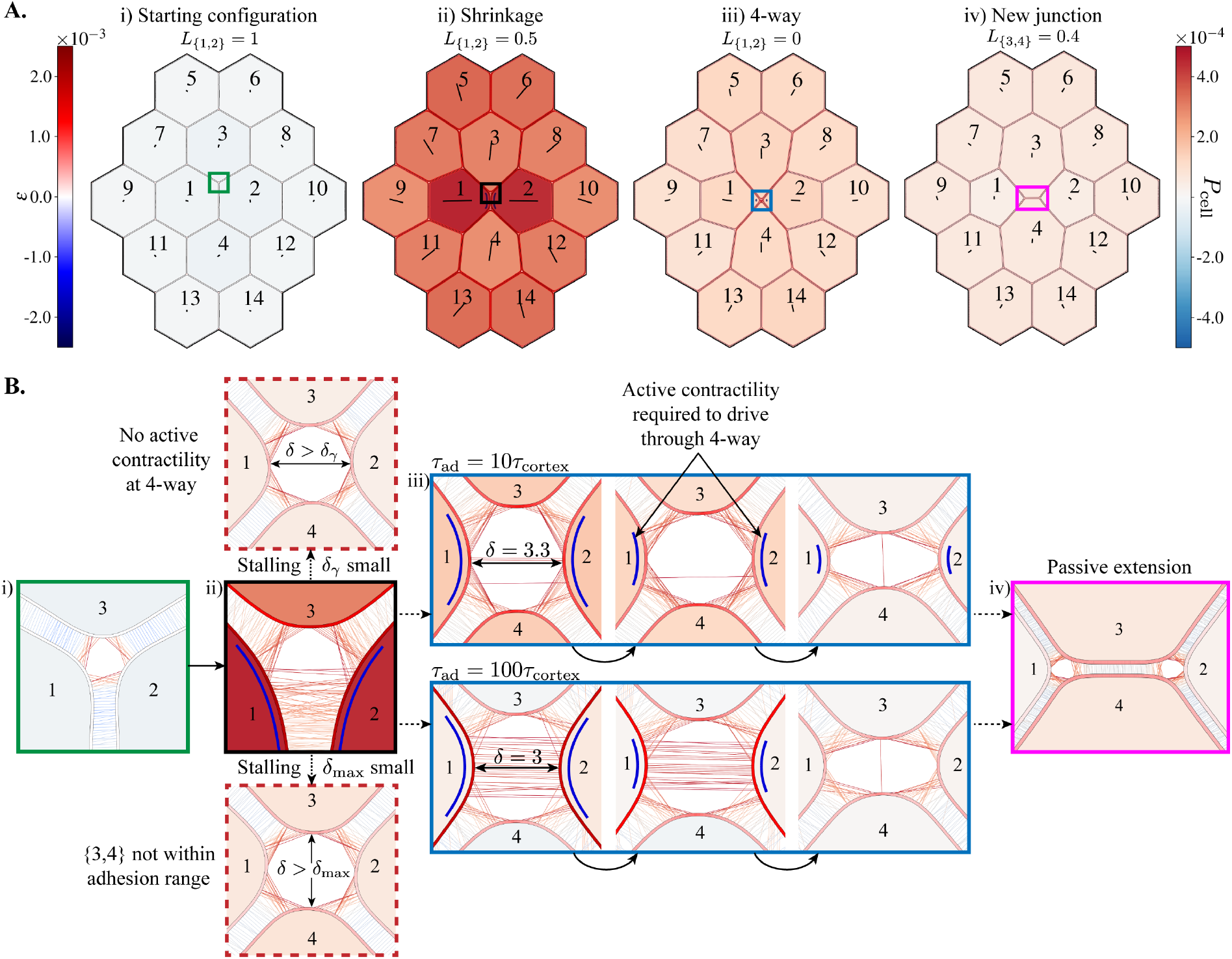
Simulating an active neighbour exchange. **(A)** Image sequence from Movie 2A, with *τ*_ad_ = 10*τ*_cortex_, (*κ, ω*) = (0.01, 0.05) and active contractility (*γ* = 0.94) in only the {1, 2} junction. All other cortex locations (in all cells) have a low-level prestress, *γ*_0_ = 1 − 2 × 10^−4^. Cell shading represents the magnitude of isotropic cell stress, *P*_cell_ (see equation (17) of the Supplementary Document), and cortex shading represents the strain, *ε* = *α* 1. Lines within cells show the principal axis of cell stress. **(B)** Close-ups of cell vertices from coloured boxes in **A**. i) Before active contractility is applied. ii) During junction shrinkage. Dashed arrows above and below show final configurations when the rearrangement is unsuccessful: the simulation stalls when the active contractility does not persist through the 4-way (top) and if cells 3 and 4 are not brought within *δ*_max_ (bottom). iii) Resolution of the 4-way vertex for two choices of the adhesion timescale. iv) Following resolution, the new junction extends passively. Cortex locations with active contractility are marked by blue lines next to the cortex.

Figure 3 shows an example simulation of an actively driven neighbour exchange (see also Movie 2A). We find that active contractility in a single junction is sufficient to shrink the junction to zero length. Importantly, however, the neighbour exchange stalls at the 4-way configuration if the active contractility is lost before new adhesion connections are made between the cells coming into contact (cells 3 and 4 in Figure 3). The model thus requires that *δ*_*γ*_ is greater than the spacing between the cells losing a junction (cells 1 and 2; Figure 3Bii, top), until the cells coming into contact are within *δ*_max_ and can form adhesion bonds (cells 3 and 4; Figure 3Bii, bottom). We highlight that using *δ*_*γ*_ to specify active contractility is a modelling choice. We could also imagine that cells 3 and 4 could be brought into contact by other means e.g. active contractility lingering in the cortex after the specification is lost, or external cell- and tissue-level forces helping to drive through the 4-way configuration.

### 2.6 Neighbour exchange is completed by passive extension of a new junction

The 4-way configuration is resolved by the appearance of a new junction, which extends passively. Then, since there are no longer adhesions between cells 1 and 2, there is no active contractility. The passive extension is due to the decreasing free energy of the system, as the bicellular junctions neighbouring the new junction form internal angles of 2*π/*3. The extension is facilitated by, but does not require, the low-level pre-stress in all cells. Moreover, the neighbour exchange and extension relax stress at the cell level, which tends to orient towards the contracting junction (see cell shading and orientation of axes in Figure 3 and Movie 2A).

Figure 3B shows the organisation of adhesion bonds through the course of the neighbour exchange, for two values of *τ*_ad_. We observe more disorganised adhesion configurations as the lifetime of adhesions increases. Larger values of *τ*_ad_ also keep the cortices in the shrinking junction more strongly coupled, reducing the inter-cortical distance (Figures 3Biii & S4). Furthermore, new adhesion bonds between the cells coming into contact (cells 3 and 4) cannot form until the existing adhesions unbind, according to *τ*_ad_. Despite this, the shrinkage, resolution and extension phases are successful for all *τ*_ad_ *< ∞* (Movie 2B), as long as active contractility endures until the 4-way is resolved. The adhesion timescale does however have important consequences for the dynamics of the process.

### 2.7 The adhesion timescale modulates cell area loss and apposed-cortex shear

Repeating the junction shrinkage simulation for several demonstrative adhesion timescale choices, we find that fast adhesion turnover (*τ*_ad_ < *τ*_cortex_) leads to significant area loss in the cells sharing the neighbouring junction (Figure 4A; compare Figure 3Aii to 4C and Movie 2A to Movie 3A). This area loss is commensurate with increased shear between apposed cortices in bicellular junctions (Figure 4B). By contrast, for slow adhesion turnover, apposed-cortex shear will gradually increase the angles between the cortex and adhesion bonds (Figure 4D). This, in turn, causes adhesion forces to oppose further shear, behaving as an effective frictional viscosity *µ*_ad_ = *ωτ*_ad_. Both the area loss and apposed-cortex shearing are dramatically reduced for any *τ*_ad_ *≥ τ*_cortex_, with area losses of *∼* 25%. Strikingly, this value is similar to the area loss observed in the cells losing a junction during convergent-extension in the *Drosophila* germ-band (Figure 4A).

**Figure 4:**
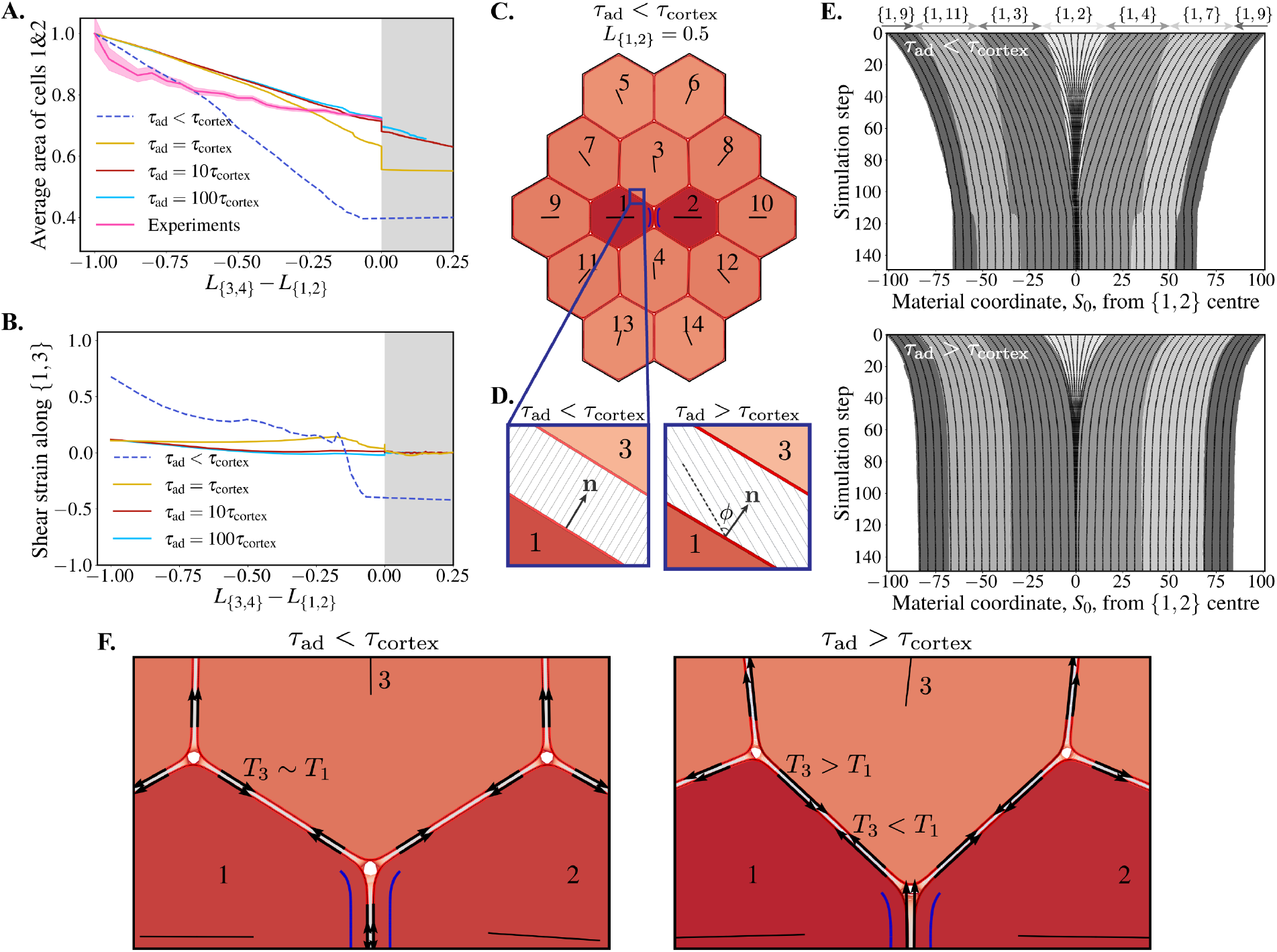
The adhesion timescale regulates area loss, shear, material flow and tension transmission. **(A)** Average area of cells 1 & 2 vs. signed length of the shrinking/growing T1 junction (*L* _{3,4}_ *− L* _{1,2}_) in simulations and experiments, normalised to their initial values. Experimental data is from wild-type cells in the *Drosophila* germ-band, reanalysed from the dataset originally published in [17]. Shading represents 95% confidence intervals. **(B)** Tangential shear strain between cortices 1 and 3 at the centre of the {1, 3} junction vs. signed length of T1 junction (as in **A**). **(C)** Snapshot from junction shrinkage simulation with fast adhesions, *τ*_ad_ < *τ*_cortex_ (see also Movie 3A), where the {1, 2} junction has had a 50% reduction in length, comparable to Figure 3Aii. **(D)** Example snapshots of adhesions along the {1, 3} junction in fast and slow adhesion timescales. On slow timescales there is a large shear angle, *φ*, relative to the cortex normal, **n. (E)** Kymographs along the entire cortex of cell 1 during junction shrinkage, for fast (top; Movie 3B) and slow (bottom; Movie 3C) adhesion timescales. Grey shading represents each bicelluar junction, with the {1, 2} junction centred at 0 and eventually disappearing. The *x*-axis shows the reference material coordinate, *S*_0_. Fixed material points (black lines; purple dots on cell 1 in Movies 3B&C) are tracked over the course of the simulation. Black lines crossing the grey shading boundaries indicates material flow between junctions. **(F)** Zoom of tissues in **C** (left; Movie 3D) and Figure 3Aii (right; Movie 3E) showing the differential tension, *T*, (black arrows) between the apposed cortices in bicelluar junctions.

### 2.8 The adhesion timescale regulates material flow across cell vertices

The larger shear rate for slower adhesion timescales is indicative of more relative material flow between junctions. As with the 3-cell tissue in Figure S2, we quantify this by tracking the locations of material points (Lagrangian tracers) in the cells. Figure 4D shows kymographs of sampled material points in the whole cortex of cell 1 over the course of a neighbour exchange. For both fast and slow adhesion turnover, we see that material points in the shrinking {1, 2} junction collapse as the junction contracts (see also blue circles on the cortex of cell 1 in Movies 3B&C). However, this effect is exacerbated when the adhesion timescale is fast relative to the timescale of the cortex, *τ*_ad_ < *τ*_cortex_, as material is dragged from the rest of the cortex into the contracting {1, 2} junction. This is an extreme case where vertices are acting as almost-frictionless pulleys, with significant cortical material flowing freely between junctions (Figure S3). In contrast, slower adhesion timescales increase the friction between apposed cortices and prevent material transfer (notice that black lines mostly stay within the same shaded grey region in Figure 4D, lower).

### 2.9 The adhesion timescale regulates tension transmission in an active tissue

Slower adhesion turnover prevents material flow across vertices, reducing shear and providing a stronger mechanical coupling between apposed cortices. We explore this further by assessing the tension in apposed cortices along bicellular junctions in different adhesion timescale regimes. When *τ*_ad_ < *τ*_cortex_, the active stress generated in the shrinking junction is held by the elasticity in the cortices of cells 1 and 2. Figure 4E (left; Movie 3D) shows that cortical tension is relatively constant around the cortex of cell 1 and is of the same order as in cortex 3. In this case, tension is transmitted around the cortex of cell 1, allowing the shrinking junction to pull material from the rest of the cortex, shrinking the cell perimeter and, therefore, the area. The stress in the neighbouring cell cortices then comes from the expansion required to fill the space thus left vacant. Conversely, when *τ*_ad_ *≥ τ*_cortex_, tension is transmitted through the lingering adhesions from cell 1 to its neighbouring cortices (e.g. cell 3 in Figure 4F, right; Movie 3E). Here, the tension in the cortex of cell 3, for example, arises from the adhesions pulling its cortical material towards the contracting junction. Note also that, as a result of these different mechanical balances, the angles formed by junctions at the vertices differ depending on *τ*_ad_.

### 2.10 The adhesion timescale defines a frictional viscosity that regulates tissue dynamics

By resisting relative tangential motion between apposed cortices, the adhesion timescale is specifying a frictional viscosity, *µ*_ad_ = *ωτ*_ad_, which modulates tension transmission and shear in the tissue. We see how this regulates the dynamics of shrinkage, resolution and extension by tracking the length of the shrinking {1, 2} junction across simulation time (Figure 5A). When adhesion dynamics are fast relative to cortex relaxation (*τ*_ad_ < *τ*_cortex_) active shrinkage is slow, since material from the rest of the cortex flows into the contracting junction and the active tension is not well-transmitted to neighboring cells (Figure 4E). However, the 4-way configuration is resolved almost immediately and followed by very fast extension, since it involves little perimeter change and the shear resistance from *µ*_ad_ is small. Shrinkage speed increases proportionally as adhesion turnover is slowed, since lingering adhesions prevent material in-flow.

**Figure 5:**
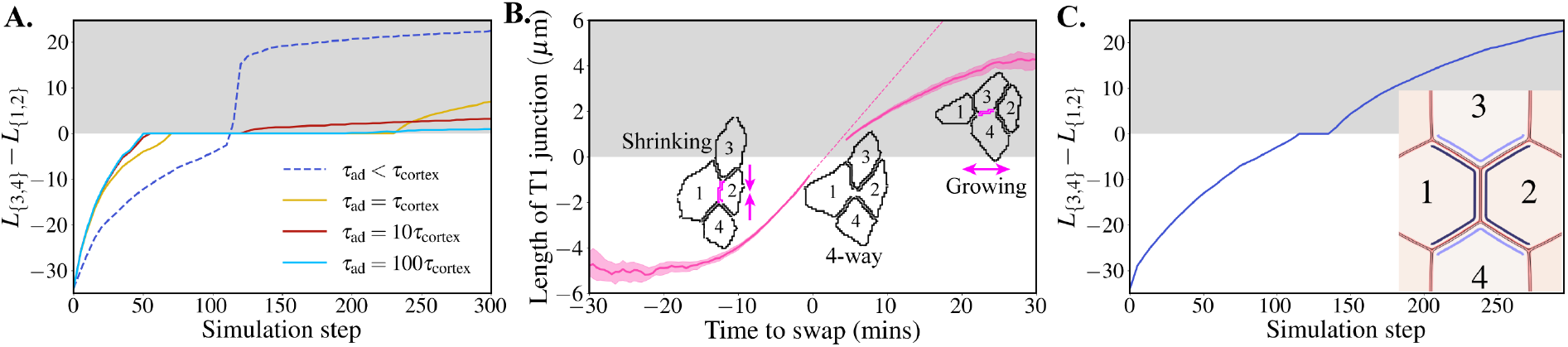
Shrinkage, resolution and extension dynamics across adhesion timescales. **(A)** Signed length of T1 junction (negative when {1, 2} shrinking; positive when {3, 4} growing, grey shading) vs. simulation step. **(B)** Experimental data of signed length of T1 junction vs. time to neighbour exchange swap for wild-type cells in the *Drosophila* germ-band. The data was extracted from experiments originally published in [23]. Lengths below 0.75 *µ*m are not plotted, due to insufficient imaging resolution. Dashed line shows continuation of shrinkage speed from 5 mins before swap, which meets re-emergence of solid line after swap, indicating a minimal delay during the resolution phase. Shading represents 95% confidence intervals. **(C)** Signed length of T1 junction vs. simulation step for a simulation where contractility is applied asymmetrically along bicellular junctions neighbouring the shrinking junction. Cortices next to darker (or lighter, resp.) blue lines in snapshot have *γ* = 0.99 (or 0.9975, resp.). See also Movie 4A.

Cells become jammed in the 4-way configuration for a length of time that is specified by the adhesion timescale and the magnitude of surrounding active stress. The adhesion timescale defines when cells 1 and 2 uncouple (unless the surrounding active stress is large enough to break the adhesions). Once uncoupled, the magnitude of active contractility defines how quickly the 4-way is resolved. Note that resolution is slowest when *τ*_ad_ = *τ*_cortex_ because, although cells 1 and 2 uncouple quickly, there is less active stress to drive through the 4-way since the cortices are further apart (see Figure S4A). This is an artefact of our implementation prescribing contractility using *δ*_*γ*_. After resolution of the 4-way configuration, tension drops dramatically. The shortest timescale now present in the system is the adhesion timescale, which therefore sets the rate of extension.

Our experimental data show that intercalating cells in the *Drosophila* germ-band resolve the 4-way configuration with almost no delay, which is remarkable (Figure 5B). We also find that junction extension is slower than shrinkage (Figure 5B). However, junction extension is proportionally faster, relative to shrinkage, than it is in our simulations. It is likely that there are multiple mechanisms facilitating extension, including anisotropic forces from within the bulk of the cells [10, 52]. We asked whether this model could reproduce the experimental shrinkage and growth rates without adding these additional forces. Indeed, we show that this can be done using a purely junctional mechanism by considering cables with asymmetric contractility within bicellular junctions. We set the cortices of cells 1, 2, 3 and 4 to be contractile when they share adhesions, creating two cables: {{1, 3}, {1, 2}, {1, 4} and {{2, 3}, {1, 2}, {2, 4}} (Figure 5C). This represents, for example, actomyosin cables that are observed in the *Drosophila* germ-band. Following experimental predictions [17], we impose that the magnitude of contractility is larger in the cortices of cells 1 and 2. Thus the bicellular junctions {1, 3}, {1, 4}, {2, 3} and {2, 4} have asymmetric contractility – a unique feature of this model. In this configuration, we successfully reduce the delay in resolution and match shrinkage speed with extension (Figure 5C and Movie 4A).

### 2.11 Large adhesion viscosity can promote rosette formation in an active tissue

The dynamic behaviour of adhesions becomes increasingly important in a highly active tissue, where multiple rearrangements may happen in succession. We simulate this case by initialising contractility in the neighbouring *{*1, 3*}* junction after the 4-way configuration has been resolved (starting from Figure 3Aiii), but before much extension of the nascent {3, 4} junction (Figure 6). When the adhesion timescale is of the order of the cortical timescale, the newly contracting junction facilitates extension of the nascent {3, 4} junction and there is a second successful neighbour exchange (Movie 5A). However, we can increase the penalty to shearing relative to contraction by slowing the adhesion timescale, increasing *µ*_ad_ (Movie 5B). In this case, resolution of the 4-way is slower than shrinkage of the newly contracting {1, 3} junction and its two vertices are dragged together without resolving the 4-way vertex first. This leads to a 5-way vertex, also known as a rosette [22, 53], being formed. Interestingly, the rosette is indefinitely stable while there is a small pre-stretch/stress, *γ*_0_, in all cells. Removing the background tension in all cells, by setting *γ*_0_ = 1 in all cells, leads to eventual resolution of the rosette. This indicates that rosettes are intrinsically unstable, but are stabilised by the low-level tension held in the cortices of the surrounding cells.

**Figure 6:**
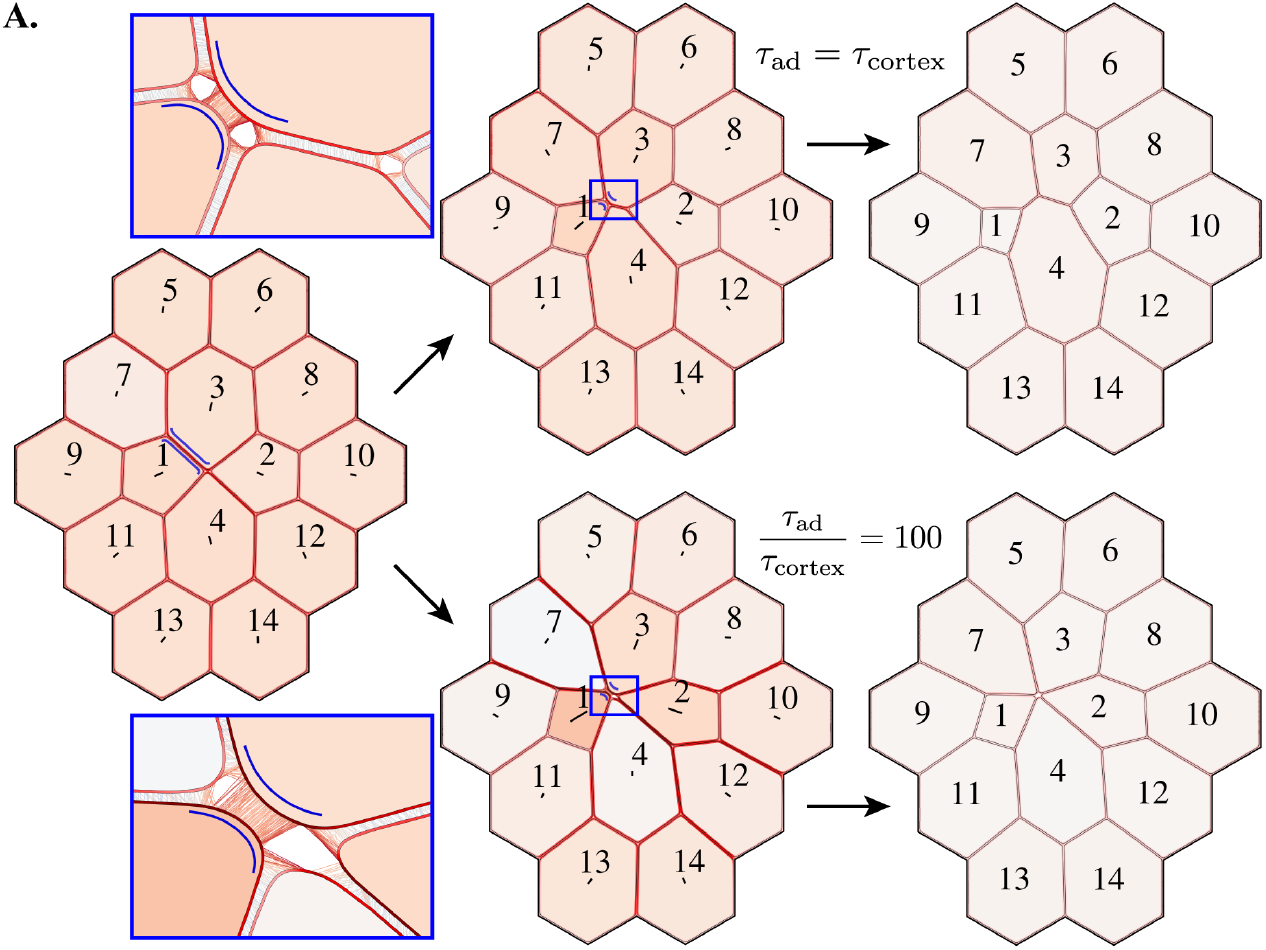
Rosette formation in a high-friction tissue. **(A)** The simulation from Figure 3 was stopped once the {3, 4} junction was formed and active contractility (*γ* = 0.94) was applied to the {1, 3} junction. If *τ*_ad_ = *τ*_cortex_ (top; Movie 5A), the {3, 4} junction extends. However, if the adhesion timescale in the tissue is slower, *τ*_ad_ = 100 *τ*_cortex_, introducing more friction to the system, the {3, 4} does not resolve (bottom; Movie 5B). Instead, two vertices are merged to form a rosette structure. The rosette is stable while all cells have a small pre-stress, *γ*_0_.

## 3 Discussion

Representing the actomyosin cortex of each cell in a tissue as an extensible morphoelastic rod, we have presented an apposed-cortex description of active epithelium dynamics. With direct access to the mechanical properties of the actin junctional cytoskeleton 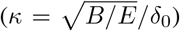, active contractility (*γ*) and adhesions (*ω*), the model has more effective control in reproducing different physiological conditions than models with a higher level of phenomenology.

We show the importance of considering whether the actomyosin cortex can flow past a vertex, from one junction to the next, and that this can be regulated by the resistance to shear between apposed cell cortices. In this model, these phenomena emerge as a consequence of – and are modulated by – adhesion turnover, whose timescale *τ*_ad_ defines a viscous friction, *µ*_ad_ = *ωτ*_ad_, that regulates the penalty to tangential shearing between apposed cortices. Relative to the cortical viscosity, *µ*_ad_ defines regimes where we can consider cortices behaving as springs pinned together at vertices versus extensible ropes passing through pulleys at vertices (Figure S3). In the latter case, junctions can change length by both contraction/expansion and by passing material past vertices. Interestingly resistance to cell area change emerges from this mechanism, rather than as a phenomenological input. These results suggest that measurements of *τ*_ad_ *in vivo*, though difficult to extract, would provide valuable insights to the emergence of different tissue behaviours.

We provide relationships between the geometric configuration of cell vertices and the mechanical properties of the cortex (*κ*) and adhesions (*ω*) that can be used to fit the model. The model predicts that cell vertices are sites of mechanical vulnerability. This is in agreement with cell culture experiments, which exhibit vertex fracturing in the presence of strong myosin II contractility [54]. Recent experimental evidence in *Drosophila* also indicates that cell vertices are mechanically unique regions in a tissue, with the localisation of vertex-specific components that both facilitate polarized cell intercalation and are important for tissue mechanics [23, 55–57]. We hypothesise that such molecules could regulate material flow past vertices. While very difficult to encode in discrete vertex-based models, the impact of vertex-specific adhesions on the biophysical properties of a tissue will be straightforward to explore in this framework.

Many existing models of epithelial mechanics, such as vertex-based models, treat cell intercalations as discrete, instantaneous events and thus cannot study the dynamics of their behaviour. We use the continuum apposed-cortex model to demonstrate that cell neighbour exchanges can be driven by purely junctional contractility. This reproduces existing experimental observations [14]. Notably, other contributions to polarized cell intercalation are known *in vivo*, such as basal protrusions first identified in vertebrate models [32, 58] and more recently in *Drosophila* germ-band extension [59]. Also in the *Drosophila* germ-band, recent studies have indicated that medial pools of myosin II, associated with the apical cell surface and distinct from the junctional pools, contribute to junction shrinkage [11]. Point forces at vertices and (possibly anisotropic) medial forces could be added to the apposed-cortex model to explore how these tissue-specific mechanisms act to facilitate junction shrinkage.

The model predicts that extension of a new junction follows passively once the 4-way configuration is resolved. This passive extension can be facilitated by a small global pre-stress. *In vivo*, additional mechanisms may facilitate the elongation process and regulate its dynamics. Indeed, in the *Drosophila* germ-band, medial pools of myosin contract asymmetrically around 4-way vertices and are thought to aid resolution and extension [10, 14, 60, 61]. Supporting this notion, we show the relative rate of junction extension can be increased by asymmetric contractility in bicellular junctions (Figure 5C). Future work should look to match the dynamics of resolution and growth to particular experimental systems to characterise the mechanical impact of supplementary mechanisms.

Rosette structures, where four or more cells share a vertex, are found in many developing tissues [20–22, 58, 62, 63]. While these structures are thought to impact the mechanical properties of tissues [64], it is often not understood why they form with higher prevalence in particular conditions [23]. We present novel predictions for how the mechanical properties of subcellular constituents can promote rosette formation and alter tissue-level structure; rosette formation is more likely in tissues in a large adhesion friction regime, where junction shrinkage (determined by cortical contractility and viscosity) happens on faster timescales than junction resolution and growth (determined by adhesion friction). Larger-scale simulations could now be used to better understand how subcellular mechanical parameters (e.g. *µ*_ad_) and tissue-scale geometric and topological properties (e.g. number of rosettes) separately contribute to tissue-level mechanical properties (e.g. shear and bulk moduli). For such large-scale simulations, future work could look to encode the rheological behaviours of this model into a more simplified description, perhaps termed a “cortex-vertex model”, which could retain the key novel effects in a computational framework of complexity similar to vertex models. Indeed, it will be interesting to understand how the mechanical parameters in this model relate to the phenomenological parameters in traditional vertex models.

## Materials and Methods

### STED imaging (Figure 1B)

To label the actomyosin cytoskeleton, we used the *Drosophila* transgenic strain *w, sqhTI-eGFP [29B]*, where the endogenous Myosin II Regulatory Light Chain is tagged with GFP [65].

Stage 8 *Drosophila* embryos from this strain were fixed and immunostained using standard procedures. The fixation step is 7 minutes in 40% formaldehyde. Primary antibodies were mouse anti-phosphotyrosine (Cell Signalling Jan-15, 1:1000) and rabbit anti-GFP (ab6556-25, abcam 1:500). Secondary antibodies were goat anti-mouse-Star red (Abberior, 2-0002-011-2) and goat anti-rabbit Star 580 (Abberior 2-0012-0050-8) (Life Technologies, 1:100).

Embryos were mounted on slides in Vectashield (Vector Labs) under a coverslip suspended by a one-layer thick coverslip bridge on either side. This flattened the embryos sufficiently so that all cells were roughly in the same z-plane. Prior to placing the coverslip, embryos were rolled so that their ventral surfaces were facing upwards towards the coverslip.

Embryos were imaged on an Abberior Expert Line STED microscope, in lateral depletion mode. Excitation was centred at 585 nm and 635 nm for Star 580 and Star Red respectively while 775 nm depletion was used for both colors. theAAn Olympus UPlanSApo 100x/1.40 Oil immersion lens was used on an Olympus inverted frame for all STED imaging. Xyz stacks were taken and a single plane is shown in Figure 1B at the position of adherens junctions.

### Data from time-lapse movies

Figure 1B shows segmented contours of cells at the level of adherens junctions, from time-lapse confocal movies of stage 7-8 *Drosophila* embryos, where intercalating cells are segmented and tracked through time (dataset originally published in [17]).

Biological data in Figure 4A is taken from the segmentation and tracking of *Drosophila* germband cells in 6 wild-type movies analysed in [17]. For each T1 event in the data (N=5580), identified when the internal interface of a quartet of neighbouring cells changes from one pair to the other (equivalent to from cell pairs 1,2 to 3,4 in Figure 3A), the length of the shortening interface and the average area of the two cells sharing the shortening interface are extracted. Mean average area is plotted against junction length, with 95% confidence intervals shown.

Biological data in Figure 5B is extracted from the segmentation and tracking of *Drosophila* germ-band cells in 5 wild-type movies, presented in [23] (and quantified there in Figure S9). Mean junction lengths (± 95% confidence intervals) are shown for all T1 events identified between 0 and 30 minutes of germ-band extension (N = 1445).

## Data Availability

The source code implementing the apposed-cortex model is publicly available for use at github.com/ Alexander-Nestor-Bergmann/appcom. Documentation and quick-start tutorials can be be found at appcom. readthedocs.io.

## Acknowledgements

The computations were performed using the Cactus cluster of the CIMENT infrastructure, supported by the Rhône-Alpes region (Grant CPER07_13 CIRA) and the authors thank Philippe Beys who manages the cluster. We thank Martin O. Lenz from the Cambridge Advanced Imaging Center for help with STED imaging. The STED microscope was funded by BBSRC grant BB/R000395/1. We thank Bruno Monier for the gift of the Sqh-GFP strain. Special thanks to Claire Lye and Alexander Erlich for their critical reading of the manuscript and members of Bénédicte Sanson’s research group for their helpful discussions. This work was supported by a Wellcome Trust Investigator Award to BS (099234Z12Z and 207553Z17Z). ANB was also supported by a University of Cambridge Herchel Smith Fund Postdoctoral Fellowship. AGF was supported by a Vice-Chancellor’s Fellowship from the University of Sheffield.

## Supplementary Figures

**Figure S1:**
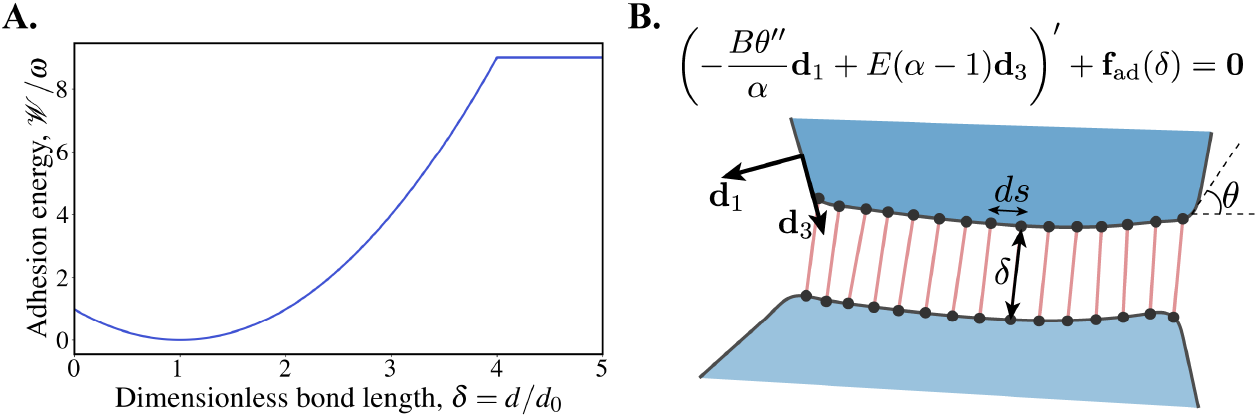
Supplementary modelling figure. **(A)** Dimensionless adhesion energy vs. bond length. The force from adhesions is modelled as a spring, with zero force at *δ* ≥ *δ*_max_ for *δ*_max_ = 4. **(B)** Numerical representation of the model. For each cortex, the bending and extensional forces balance the forces from adhesion. Adhesions act as springs, of length *δ*, acting over discretised segments *ds*.

**Figure S2:**
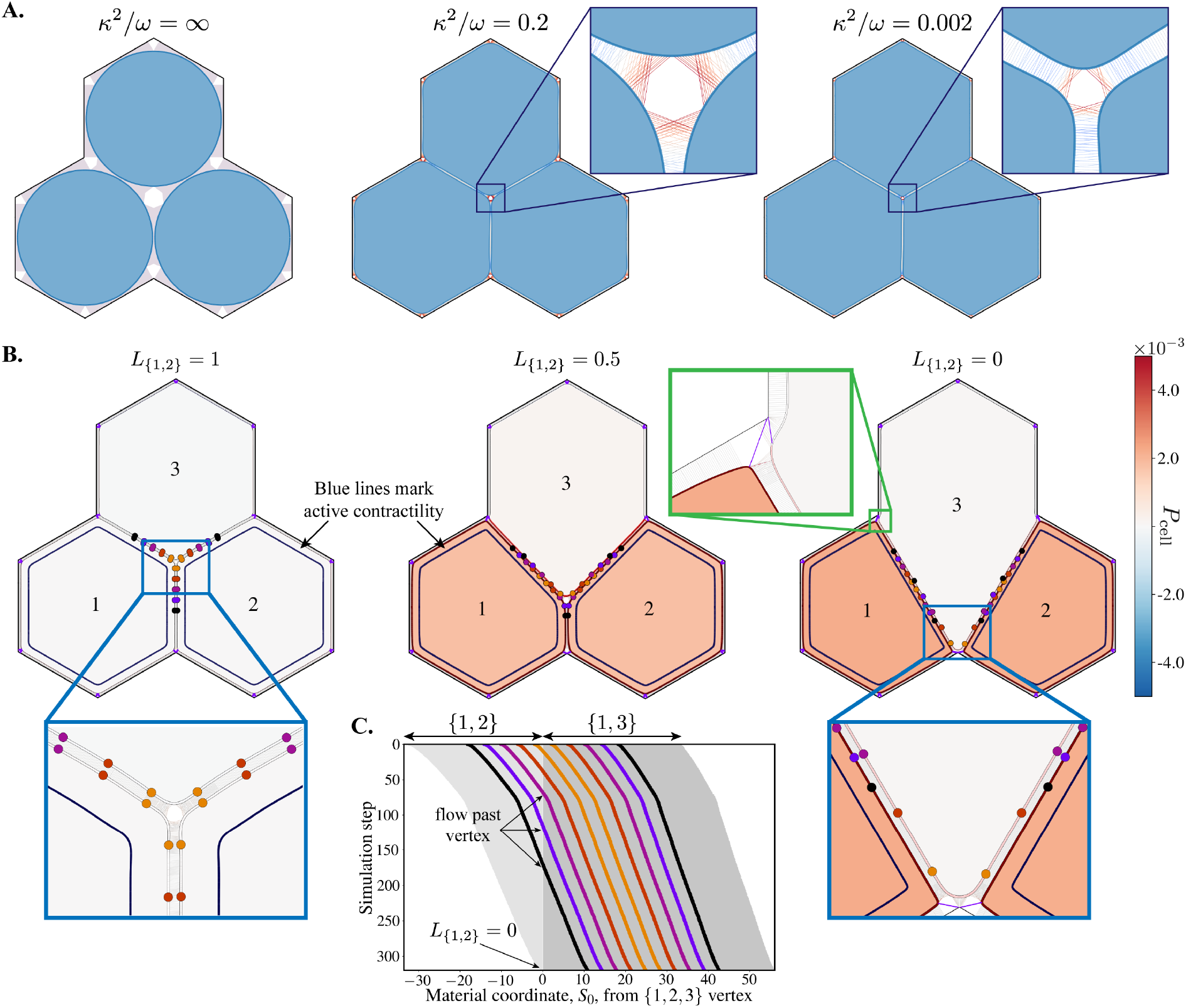
Active mechanics affect cell vertex stability and shearing behaviour. **(A)** Representative 3-cell tissues across parameter space. The right-most example is the *κ*^2^*/ω* = 0.002 isoline with coloured circles in Figure 2B. **(B)** Snapshots from Movie 2B, driving junction shrinkage with asymmetric contractility. The whole cortices of cells 1 and 2 are contractile (dark blue line in cells shows where active contractility has been applied). Coloured dots represent fixed material (Lagrangian) points that flow past one another, between apposed cortices, demonstrating cell-cell shear. Boundary vertices have been pinned with extra-stiff adhesions (50*ω*; purple lines highlighted in green box) to maintain vertices at boundary angular points. Cell shading represents the magnitude of isotropic cell stress *P*_cell_. **(C)** Kymograph along junctions {1, 2} and {1, 3} in the cortex of cell 1, showing the motion of material points (coloured Lagrangian markers) during the simulation shown in **B** and Movie 1B. The *x*-axis origin is at the transition from the {1, 2} to {1, 3} junction. Grey shading represents the extent of each junction, with darker grey showing growing {1, 3} and lighter showing the shrinking {1, 2}. Coloured lines crossing the boundary between light/dark grey shading indicate material points flowing across the vertex, from {1, 3} to {1, 2}.

**Figure S3:**
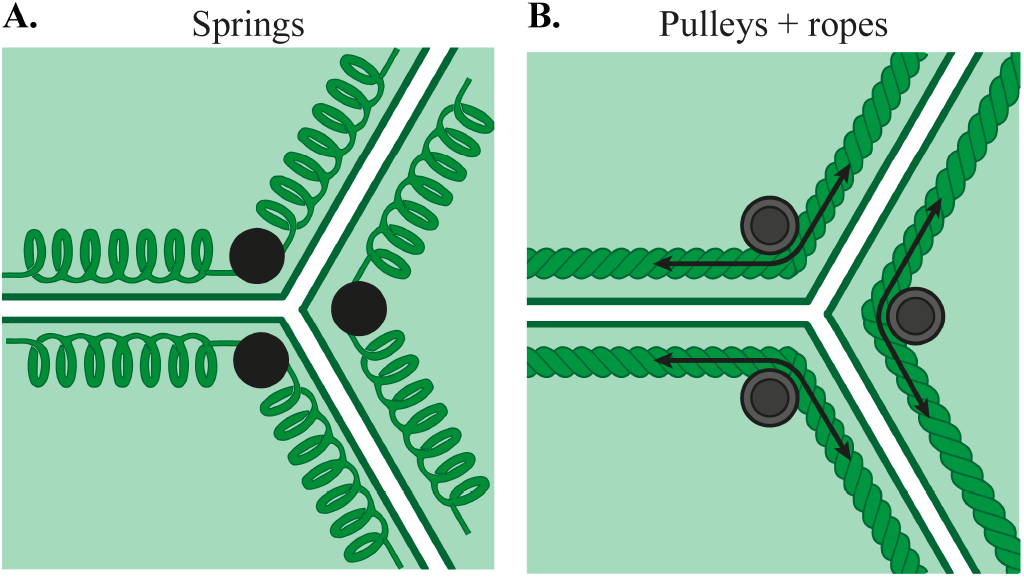
Mathematical representations summarising junction mechanics: pinned springs or ropes and pulleys? **(A)** A common constitutive model of the cortex, where cell junctions behave as springs pinned at vertices. **(B)** A plausible alternative representation of the cortex, where material can be passed between junctions. Cortices behave as extensible ropes that can flow around a pulley at the vertex. In the apposed-cortex model, presented in this manuscript, the friction associated with flow between junctions (material moving across the pulley) comes from the properties of cell-cell adhesions, which anchor cortices and define a frictional shear viscosity.

**Figure S4:**
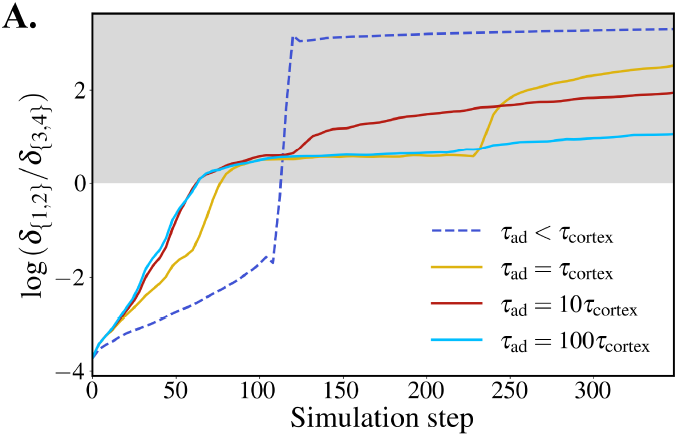
Apposed-cortex spacing anisotropy during neighbour exchange. **(A)** Log ratio of the minimum distance between cells 1 and 2, *δ* _{1,2}_, over the minimum distance between cells 3 and 4, *δ* _{3,4}_, against simulation time. The ratio being zero implies that the cells are equidistant and the 4-way vertex is isotropic in shape.

## Legends for Movies

**Movie 1A (associated with Figure 2C)**. A simulation of a tissue with three cells, where active contractility is increasing in the whole cortex of all cells (*γ* decreasing from 1) until the tissue fractures. Cell shading represents the magnitude of isotropic cell stress, *P*_cell_, and cortex shading represents the strain, *ε* = *α −* 1.

**Movie 1B (associated with Figure S2B)**. A simulation of a tissue with three cells, where junction shrinkage is driven by asymmetric active contractility. There is active contractility (*γ* = 1 − 0.005) in only cells 1 and 2. Cell shading represents the magnitude of isotropic cell stress, *P*_cell_, and cortex shading represents the strain, *ε* = *α −* 1.

**Movie 2A (associated with Figure 3)**. A simulation where junction shrinkage is driven by active contractility in a single bicellular junction. There is active contractility (*γ* = 1 − 0.04) in the junction shared between cells 1 and 2 (contractility is prescribed where the two cortices are within range *δ*_*γ*_ = 4) and *τ*_ad_ = 10*τ*_cortex_. Cell shading represents the magnitude of isotropic cell stress, *P*_cell_, and cortex shading represents the strain, *ε* = *α −* 1.

**Movie 2B**. A junction shrinkage simulation with identical active contractility conditions to Movie 2A, applied to the {1, 2} junction. The adhesion timescale was set to infinity, *τ*_ad_ = *∞*, such that adhesions do not unbind except where cortex mesh nodes are adaptively removed. Cell shading represents the magnitude of isotropic cell stress, *P*_cell_, and cortex shading represents the strain, *ε* = *α −* 1.

**Movie 3A (associated with Figure 4C)**. A junction shrinkage simulation with identical active contractility conditions to Movie 2A, applied to the {1, 2} junction. A fast adhesion timescale was used, *τ*_ad_ < *τ*_cortex_, such that adhesion bonds do not persist for multiple timesteps and instead exert a mean-field force (see equation (22) in the Supplementary Document). Cell shading represents the magnitude of isotropic cell stress, *P*_cell_, and cortex shading represents the strain, *ε* = *α −* 1.

**Movie 3B (associated with Figure 4E, top)**. An alternative visualisation of the simulation in Movie 3A, where *τ*_ad_ < *τ*_cortex_. In this movie, a sample of fixed material points are tracked on cortices of cells 1,2,3 and 4. We see that material in cortices 1 and 2 flows into the shrinking junction, indicating shearing behaviour between neighbouring cells.

**Movie 3C (associated with Figure 4E, bottom)**. An alternative visualisation of the simulation in Movie 2A, where *τ*_ad_ = 10*τ*_cortex_. In this movie, a sample of fixed material points are tracked on cortices of cells 1,2,3 and 4. We see that cortical material in cells 1 and 2 does not tend to flow between junction, across vertices, compared to Movie 3B.

**Movie 3D (associated with Figure 4F, left)**. A zoom, around the {1, 2} junction, of the simulation shown in Movie 3A, where *τ*_ad_ < *τ*_cortex_. Black arrows show the magnitude of tension in cell junctions. We see that tension roughly equal across all junctions as the tension from the contracting {1, 2} junction is held by the whole cortices of cells 1 and 2.

**Movie 3E (associated with Figure 4F, right)**. A zoom, around the {1, 2} junction, of the simulation shown in Movie 2A, where *τ*_ad_ = 10*τ*_cortex_. Black arrows show the magnitude of tension in cell junctions. We see that tension from the contracting {1, 2} junction is transmitted through adhesion bonds into the cortices of neighbouring cells.

**Movie 4A (associated with Figure 5C)**. A simulation where two contractile cables are formed in the tissue: {{1, 3}, {1, 2}, {1, 4} and {2, 3}, {1, 2}, {2, 4} (see dark blue lines next to cortices). However, the cables are asymmetric along bicellular junctions, with the active contractility being stronger in the cortices of cells 1 and 2 relative to cells 3 and 4; cortices next to darker (or lighter, resp.) blue lines have *γ* = 0.99 (or 0.9975, resp.). Cell shading represents the magnitude of isotropic cell stress, *P*_cell_, and cortex shading represents the strain, *ε* = *α −* 1.

**Movie 5A (associated with Figure 6)**. Multiple neighbour exchanges in a tissue where the adhesion timescale is comparable to the cortical relaxation timescale, *τ*_ad_ = *τ*_cortex_. Following removal of the {1, 2} junction and resolution of the 4-way vertex (as in Movie 2A), active contractility is engaged on the {1, 3} bicellular junction. This leads to a second neighbour exchange, with formation of a new junction between cells 4 and 7. Cell shading represents the magnitude of isotropic cell stress, *P*_cell_, and cortex shading represents the strain, *ε* = *α −* 1.

**Movie 5B (associated with Figure 6)**. Rosette formation in a tissue with large adhesion viscosity, *τ*_ad_ = 100*τ*_cortex_. Following removal of the {1, 2} junction and resolution of the 4-way vertex (as in Movie 2A), active contractility is engaged on the {1, 3} bicellular junction. This leads to the formation of a rosette structure, where a vertex is shared between 5 cells. Cell shading represents the magnitude of isotropic cell stress, *P*_cell_, and cortex shading represents the strain, *ε* = *α −* 1.

## Supplementary Document

This document outlines a mechanical model of an epithelial tissue with active material properties. In a departure from many previous models, we represent the junctional cortices of individual cells explicitly, with adhesion forces acting between the cortices of neighbouring cells.

### Geometry and kinematics

The cell cortex is modelled as a morphoelastic rod. We consider three configurations for the cortex (Figure 1D):

- the *initial* configuration, *𝒞*_0_, parameterised by a Lagrangian coordinate, *S*_0_ *ϵ* [0, *L*_0_], representing the arc length. This is the configuration that is adopted by the cortex at time *t* = 0, where the cortex is undeformed and has no active stress.
- The *reference* (virtual) configuration, *𝒱*, parameterised by *S ϵ* [0, *L*]. This is the unstressed configuration adopted by the cortex at time *t*. It is a conceptual configuration that the cortex adopts in response to active (pre-) stresses, but in the absence of external forces and boundary conditions. Due to physical constraints, the reference configuration may not be physically realisable in Euclidean space and can be defined only locally. For example, active stresses in the cortex may lead to a self-intersecting geometry, but this would be prevented in physical space.
- The *current* configuration, *𝒞*, parameterised by *sϵ* [0, *l*]. This is the actual configuration that is adopted by the cortex in physical space, balancing pre-stresses, body loads and boundary conditions.

These configurations are related through a multiplicative decomposition. Material elements in the cortex are taken from the initial, undeformed configuration, *C*_0_, to the reference (virtual) configuration, *𝒱*, by an active stretch:

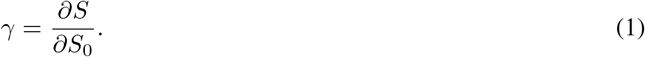

The active stretch represents local changes in material length, with *γ >* 1 for active growth and *γ <* 1 representing active contraction. Setting *γ <* 1 represents active contractility in the cortex, for example modelling the effect of bound myosin motors on the actin cytoskeleton. The mechanical effect of *γ* is to impose a residual stress, or pre-stress, in the cortex. Elastic stresses are generated when the cortex subjected to external loading and boundary conditions, taking material elements from the virtual, *𝒱*, configuration to current, *𝒞*, configuration via an elastic stretch:

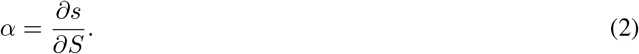

The total stretch, *λ*, between the initial and current configuration then satisfies

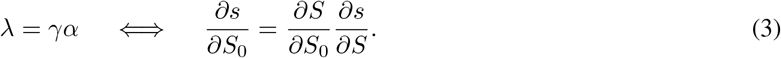

This decomposition is illustrated in Figure 1D.

In modelling the cortex as a morphoelastic rod, we follow the notation [1] and Chapter 6 in particular. Starting from the full 3D description, we provide simplified 2D equations describing the mechanics of the cortex in the apical plane. Under the rod representation, the cell cortex is defined geometrically by the position of the centre line, **r**(*S, t*), relative to the reference frame, mapping *S* to the fixed Cartesian basis {**e**_1_, **e**_2_, **e**_3}_ via **r** = *x***e**_1_ + *y***e**_2_ + *z***e**_3_. In the general frame, the rod is characterised by an orthonormal basis, {**d**_1_, **d**_2_, **d**_3}_. We set **d**_3_ to be aligned with the tangent of the centre line, normal to the rod cross section, and **d**_1_ and **d**_2_ are chosen to lie in the principal directions of the cross section, with **d**_3_ = **d**_1_ *×* **d**_2_. The full kinematic description of the cortex is given by

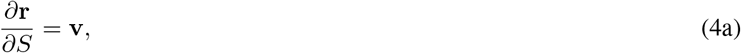

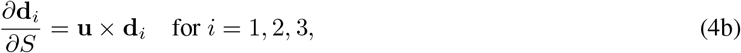

where **v** is a stretch vector (|**v**| *>* 0) and **u** is the Darboux vector. We detail these quantities under our specific assumptions below.

We simplify the generalised three-dimensional formulation with some key assumptions. Since epithelial dynamics are often driven in the apical plane, we model only the apical cortex and assume that it remains planar (torsion free). Furthermore, we assume that the cortex is unshearable, has a circular cross section and is naturally straight (no reference curvature). Importantly, however, the cortex is extensible. Under these conditions, **d**_3_ = cos *θ***e**_1_ + sin *θ***e**_2_ represents the rod tangent, **d**_1_ is the rod normal and **d**_2_ = **e**_*z*_. The angle, *θ*(*S*), is the deflection of the cortex relative to the **e**_1_ axis. From these conditions, we have [1]

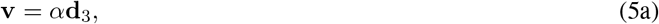

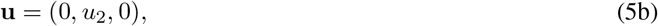

where 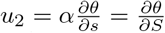 is related to the Frenet curvature, 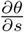, of the cortex. Thus, the full kinematic description (4) is simplified to

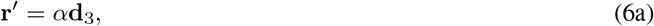

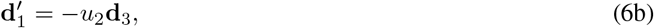

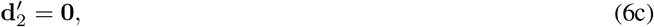

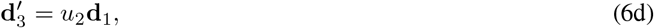

where derivatives w.r.t. *S* (virtual) are denoted (·)^*′*^.

### Cortex mechanics

With respect to the reference (virtual) configuration the balance of linear and angular momentum in the cortex give

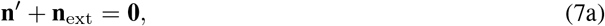

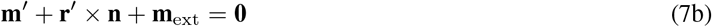

where **n**(*S*) is the force in a cross-section of the cortex, **n**_ext_ is the external force, **m**(*S*) is the moment and **m**_ext_ is the external moment. Following our assumptions above, the cortex behaves as a planar rod, extensible in the tangential **d**_3_ direction only, such that the force can be written

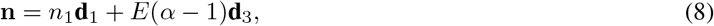

where *E* is the extensional modulus of the cortex and *n*_1_ is unknown. The cortex moment is reduced to

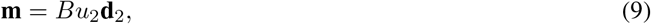

where *B* is the bending modulus of the cortex. Using (8) and (9) with (7b), assuming no external moment, we find the component of force normal to the cortex

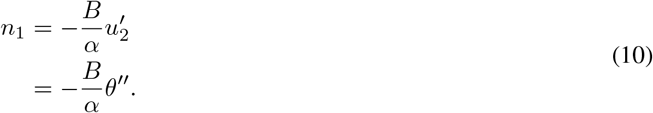

Then, the balance of linear momentum (7a) gives

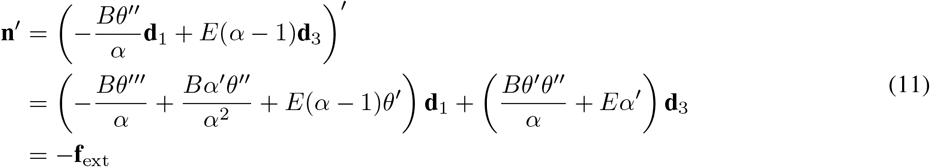

Thus we have

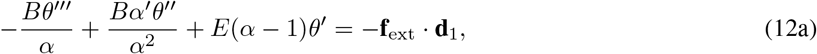

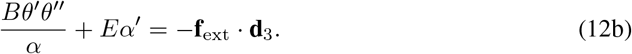

These equations describe the position of the cortex in the deformed configuration relative to the reference (virtual) configuration. However, for our purposes, it is more convenient to work with reference to the initial, undeformed configuration, upon which we can impose the active pre-stresses. We can make the change of variables, writing

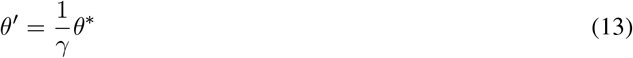

and

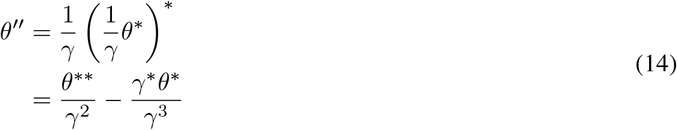

where (.)^*\**^ = *∂*(.)*/ ∂S*_0_. With respect to the undeformed configuration, we obtain a sixth-order system with four unknowns (*θ, α, x, y*):

Find (*θ, α, x, y*) : *s ϵ* [0, *l*] *→* ℝ^4^ for which

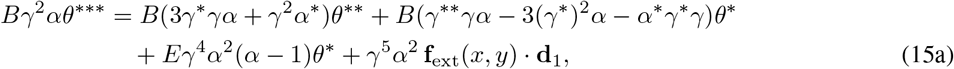

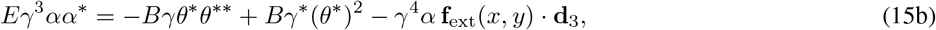

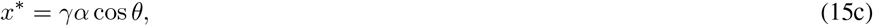

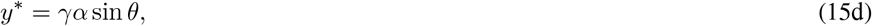

where (15c) and (15d) provide the position of the cortex in the current configuration. The system (15) is closed with the following periodic end-point conditions: (*θ*^*\* \**^, *θ*^*\**^, *θ, α, x, y*)|_*s*=*l*_ = (*θ*^*\* \**^, *θ*^*\**^, *θ, α, x, y*)|_*s*=0_ + (0, 0, 2*π*, 0, 0, 0).

#### Cell-level stress

We define the stress tensor, ***σ***, over the region, *𝒜*, of the cell enclosed by the cell boundary, *𝓁*. In the current configuration, the stress tensor is symmetric and divergence free in equilibrium, satisfying ***σ*** = ∇ · (**r** *⊗* ***σ***). Taking an integral over the cell area and applying the divergence theorem we have [2, 3]

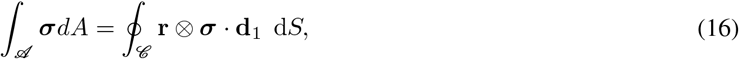

such that the cell-level stress tensor, averaged over the cell area, *A*, can be evaluated as

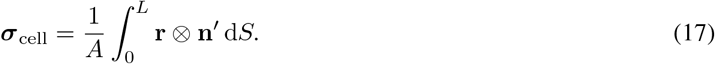

The symmetry of stress is assured by the moment balance. The trace of the stress tensor dictates the sign of its larger eigenvalue. We can therefore use the total cell pressure, *P*_cell_ = −tr(***σ***_cell_)*/*2, to characterise the state of stress within the cell: cells with *P*_cell_ *>* 0 (*P*_cell_ *<* 0) are under net tension (compression), with the principal axis of cell stress being tensile (compressive).

#### Timescales and cortical viscosity

The cortex relaxation timescale, *τ*_cortex_ *∼* 50 s, is largely driven by the timescale of actin turnover [4, 5]. This timescale is much shorter than the timescale over which rearrangement occurs (*∼* 20 min [6]). We therefore assume that the cortex behaves as a purely fluid material and update the rest length to the current length at every time step. Dynamics in the system can then be driven by changes in the mechanical properties of cells (e.g. pre-strain, rest length and stiffness) or the boundary conditions.

Many discrete models, such as vertex-based models, impose friction directly on cell vertices or other discrete material points [7–9]. This is traditionally imposed as a dissipative drag with the substrate, though attempts have been made to link dissipation to changes in cell lengths and areas [10, 11]. We extend this within the apposed-cortex framework to account for both dissipation in the cell cortex, on timescales *τ*_cortex_, as well as friction/drag between apposed cortices due to adhesion connections, on timescales of *τ*_ad_ (discussed below). We neglect inertia and dissipative interactions with neighbouring fluids and structures further apically and basally”/”in the direction *d*_2_.

### External forces: cell-cell adhesions

We consider the nature of the distributed external body forces, **f**_ext_, acting on the cortex whose work must balance the variation of mechanical energy in the cortex at equilibrium.

Adhesions are a key mechanical feature that regulate tissue stability and are a primary source of external forces acting on cell cortices. We model adhesion complexes at an effective level, imposing forces that may be a composite of contributions from E-cadherin, *α*- and *β*-catenin, vinculin and other molecules linked to the adhesion complex. The adhesion complex, as a whole, is modelled as a Hookean spring, of rest length *δ*_0_, with a maximum binding length *δ*_max_. Numerically, we represent each adhesion as a single spring-like bond at a discretisation point, weighting its force by the bond density around that point. Sufficiently far from cell vertices, the cortices of neighbouring cells are straight and parallel, and the spacing between them in the current configuration is *δ*_0_. Let us consider two neighbouring cells, *i* and *j*. At every position of coordinate *s*_*i*_ on the cortex of cell *i*, an adhesion may connect it to an available binding locations on cortex *j*, at coordinate *s*_*j*_, within a distance *δ*_max_. The line force density from a single connection between *s*_*i*_ and *s*_*j*_ is

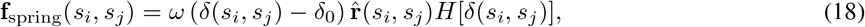

where *ω* is the product of the stiffness of a single adhesion bond and the line density of bonds, *δ*(*s*_*i*_, *s*_*j*_) = |**r**(*s*_*j*_) − **r**(*s*_*i*_)| is the distance between cortex coordinates *s*_*i*_ and *s*_*j*_ and 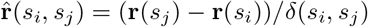 is a unit vector pointing from *s*_*i*_ to *s*_*j*_. *H*[*δ*(*s*_*i*_, *s*_*j*_)] is a Heaviside step function, defined as

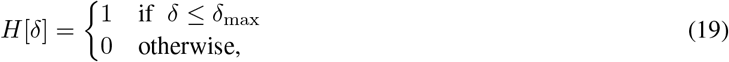

which removes force contributions from adhesion connections at a distance greater than *δ*_max_. We see that the adhesion bonds strongly couple the equilibrium equations of neighbouring cells (12).

#### Adhesion timescales

FRAP experiments have revealed that adhesion recovery times are fast (*∼* 20 s [12, 13]) compared with the timescales of cortical relaxation and neighbour exchanges. We therefore impose that adhesion binding times in the model are fast, such that unbound cortex locations form new bonds instantaneously whenever there is a neighbouring cortex within *δ*_max_. We allow for the possibility of connection to a node that already has another adhesion.

In addition to recovery time, there is an additional – less considered – adhesion timescale: the average lifetime of an adhesion complex, *τ*_ad_. This timescale represents the average time taken from binding to unbinding. It is notably distinct from, and more difficult to measure than, the adhesion recovery time. To our knowledge, there have been no explicit measurements of this additional timescale. However, the aggregated force from adhesions acting on the cortex will be strongly affected by it. We can consider it relative to the timescale of the cortex, *τ*_cortex_, and model separately the cases for *τ*_ad_ ≪ *τ*_cortex_ and *τ*_ad_ *≥ τ*_cortex_ below.

#### Slow adhesions

*τ*_ad_ *≥ τ*_cortex_. An adhesion bond pairing position *s*_*i*_ on cortex *i* to *s*_*j*_ on cortex *j* exerts a force

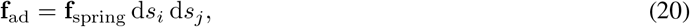

where the force is averaged over the segment lengths, d*s*_*i*_ and d*s*_*j*_. The total force acting at *s*_*i*_ on cortex *i* is then the sum of adhesion force to all points *s*_*j*_, *A*(*s*_*i*_), connected to that location :

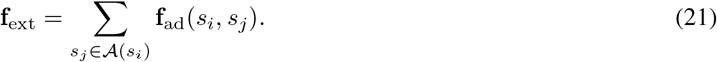

In the case where the adhesion timescale is slower than the cortical timescale, adhesion bonds persist and maintain connections between neighbouring cortices between successive iterations of simulated cortex equilibria. In this case, all existing adhesion bonds in the tissue, and their cortex-cortex pairs (*s*_*i*_, *s*_*j*_), must therefore be tracked. For simplicity, we assume that unbound cortex locations form new bonds to the nearest available neighbouring cortex.

#### Fast adhesions

*τ*_ad_ ≪ *τ*_cortex_. When the adhesion timescale is significantly shorter than the cortical timescale, adhesion forces are averaged to work in the mean-field. In this regime, individual adhesion bonds need not be tracked. Instead, the cumulative adhesion force acting on the cortex is calculated based on the position of neighbouring cortices. Considering a tissue comprising *N*_*C*_ cells, labelled *i* = 1, …, *N*_*C*_, we define the total adhesion force at cortex coordinate *s*_*i*_, on cell *i*, as the deterministic mean-field from all possible connections to neighbouring cortices within *δ*_max_:

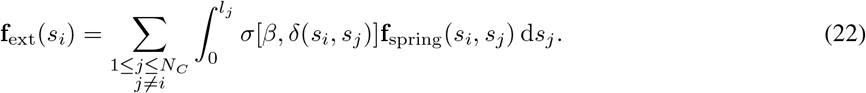

The sum is over all other cells in the tissue, but the Heaviside in (18) ensures that force contributions are taken only from cortices within *δ*_max_. Furthermore, *σ*[*β, δ*] ∈ [0, 1] is an exponential normalisation function given by

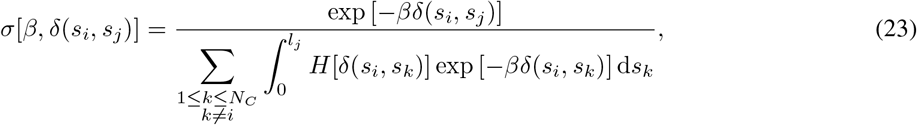

which takes the exponential ratio of the length of a single adhesion over the sum of the lengths of all adhesions connected to *s*_*i*_. The normalisation function thereby uses the length of each possible adhesion to generate a probability of connection, which is used as a scaling factor to decrease the contribution from connections to cortices that are far away. The parameter *β >* 0 controls the bias of the function, where larger values produce a stronger bias to adhesions that are nearby. In the limit *β → ∞* the total force, **f**_ext_(*s*_*i*_), is given by the force from the nearest adhesion connection. For *β* = 1 we have the conventional softmax function in base *e* and for *β* = 0 we have a uniform distribution. However, we suggest a value of *β ≥* 10, as smaller values do not provide enough bias, leading to nearby adhesions having the weakest contribution to the total force. Figure 1C provides an example of the possible adhesions connected to a point on the cortex, at a vertex, and the scaled force produced by the adhesions under (22).

### Nondimensional governing equations

We nondimensionalise lengths on the adhesion rest length (equivalent to the bicellular spacing), *δ*_0_, and use

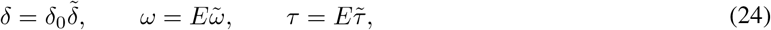

using tildes to denote dimensionless variables. The dimensionless force balance in the normal and tangential directions (per unit *S*) is then

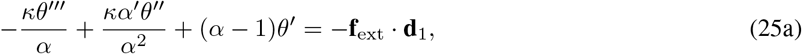

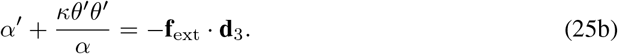

where 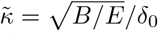 is a nondimensional parameter corresponding to the relative length scale over which bending affects the cortex around the vertex. The dimensionless force from a single adhesion bond is

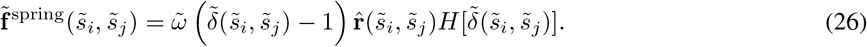

Solving with respect to the deformed configuration, we can define a dimensionless state vector of unknowns 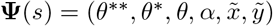. From (15), we are required to solve:

Find 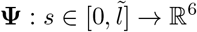 such that

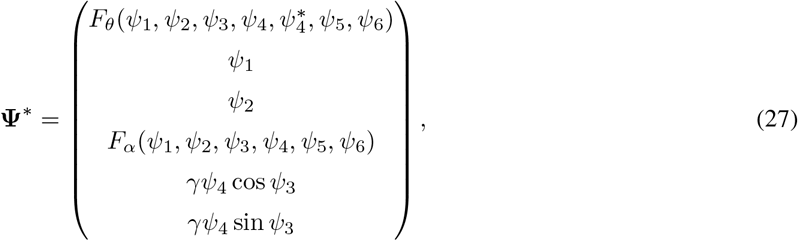

with

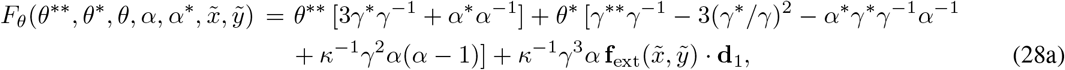

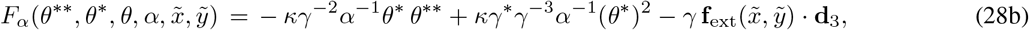

where **d**_3_ = cos *θ***e**_1_ + sin *θ***e**_2_ and **d**_1_ = − sin *θ***e**_1_ + cos *θ***e**_2_. The system is subject to periodic boundary conditions 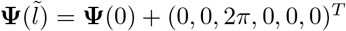.

### Parameter selection

When initialising a cell, we must set a dimensionless reference length, relative to the spacing between apposed cortices (set to 1 in the nondimensionalisation). In the *Drosophila* germ-band, cells have a radius of approximately 3.5 *µ*m prior to gastrulation [6, 14]. Conversely, cells in the developing wing disc epithelium have a smaller radius of *∼* 1 − 3 *µ*m (based on area measurements of 5 − 30 *µ*m^2^ [15]). Electron microscopy has found that the spacing between apposed cell cortices is approximately 30 − 40 nm (estimated using scale bar in Figure 7 of [16]). These observations put the ratio of the cell radius to the bicellular spacing in the range 25 − 110. Cells with larger radii are more computationally expensive to simulate. We therefore work in the lower bound and initialise cells (as a circle) with a dimensionless radius of 35, as a representative value.

It is difficult to measure the cortex extensional and bending moduli and the stiffness of adhesion bonds *in vivo*. Figure 2B of the main text demonstrates that the ratio of the dimensionless parameters, *κ*^2^*/ω*, can be fitted to the size of openings around cell vertices. To our knowledge, due to difficulties in achieving the required resolution, these measurements have not yet been reported. Using STEAD microscopy, we do not observe significant openings around cell vertices. We therefore choose order-of-magnitude estimates *κ* = 1 × 10^−2^, *ω* = 1 × 10^−2^, which give *δ*^vert^ *∼* 1.33, for this representative study.

### Deriving the mechanical balance from the energetic formulation

In the main text, we present the model in terms of the mechanical energy (returning to the dimensional model):

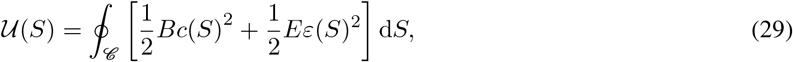

where *c* = *θ*^*′*^. We demonstrate that this is equivalent to the rod formulation described above by deriving the balance of linear and angular momentum in the cortex from **U**. We follow a variational approach, considering an infinitesimal perturbation from an arbitrary configuration of the cortex. The first variation of the mechanical energy gives the forces and torques in the cortex and at the boundaries:

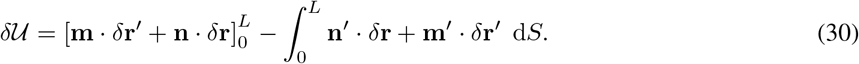

We derive these terms by taking the first variation of (29) to get

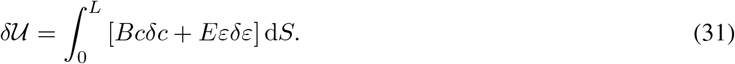

Let us consider the bending and extensional terms separately. For the bending term, note that 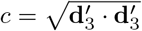 such that

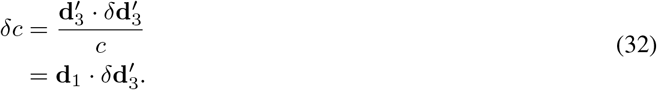

Performing two rounds of integration by parts, we have

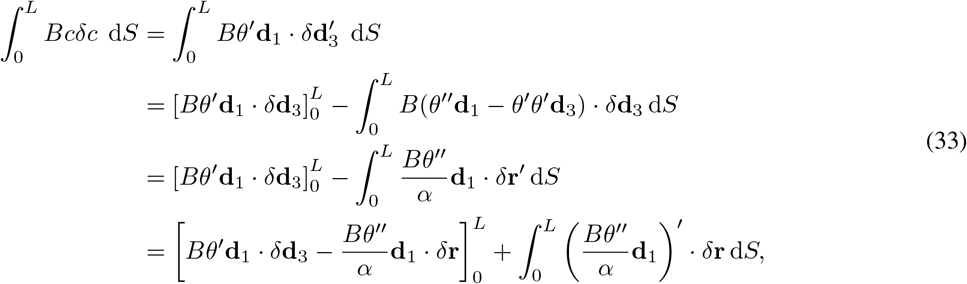

where we have used the torsion-free kinematic identities (6) and the fact that *αδ***d**_3_ = *δ***r**^*′*^ − (**d**_3_ *· δ***r**^*′*^)**d**_3_. For the extensional term, we note that 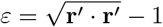 such that

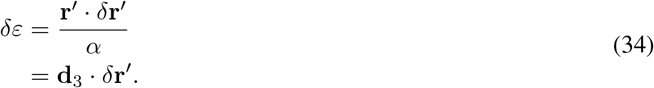

Again, using integration by parts we have

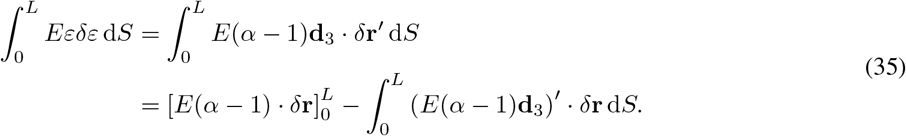

Collecting terms in (33) and (35), referencing (30), we have no net moment and the force gradient in the cortex is given by

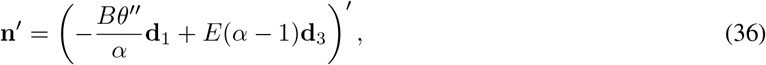

which is equivalent to (11). The terms evaluated at the boundaries impose moment and force continuity at the end-points.

### Numerical implementation and pseudocode

A pseudocode implementation of the model is provided in Algorithm 1.

**Algorithm 1:** Apposed-cortex model pseudocode

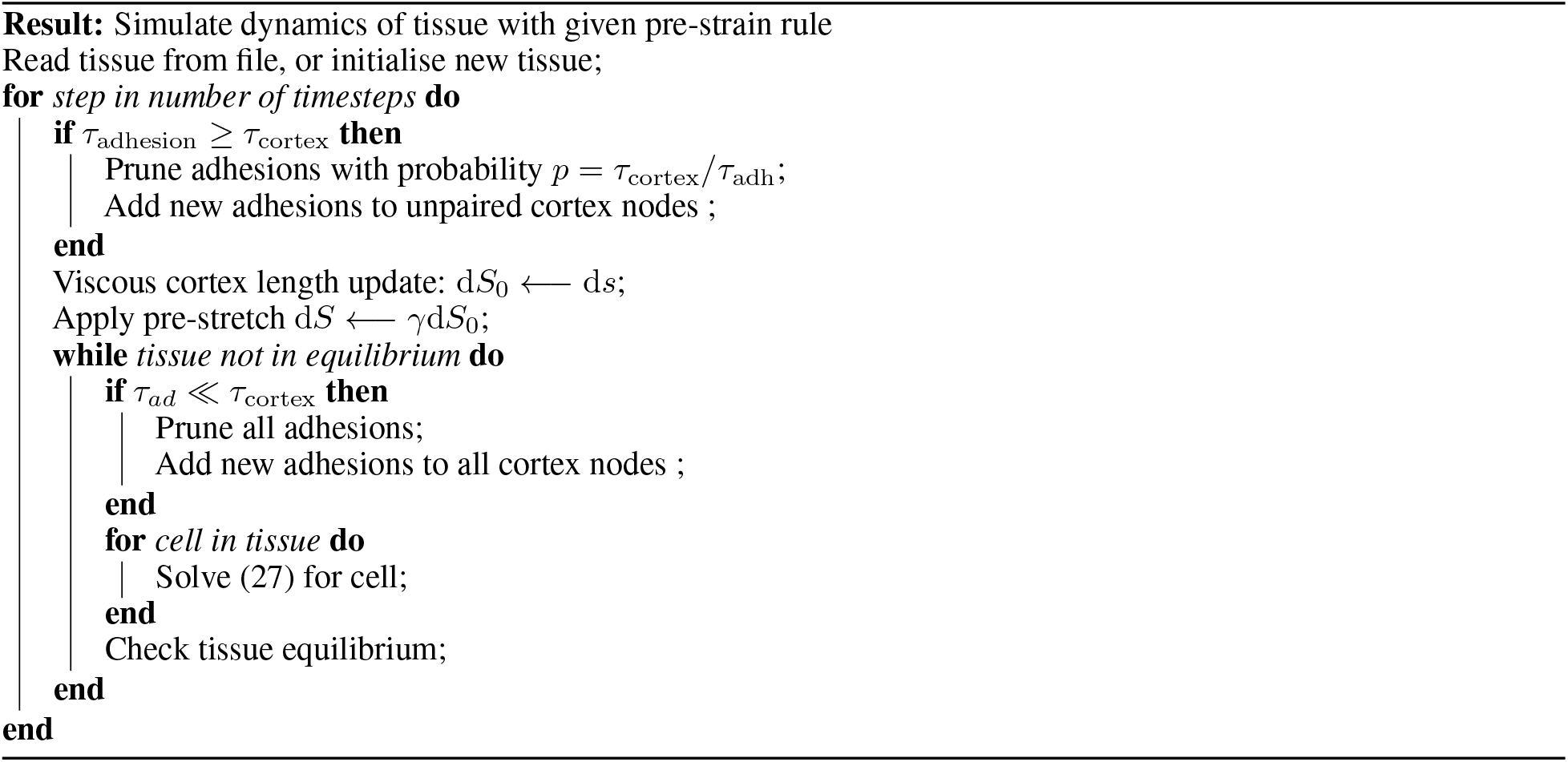

The initialisation of a new tissue can be done as described in the main text, by duplicating a single cell fitted within a hexagon. Alternatively, it is also possible to initialise multiple circles at randomised locations within a global stencil and relax them all simultaneously. However, this requires smaller changes in the adhesion strength since neighbouring cells move simultaneously and must be prevented from overlapping, thus it requires increased computational time and can be less numerically stable.

Simulations were performed in Python 3. The system (27) is solved using the solve_bvp function from the Scipy library [17]. The function performs discretisation using a fourth-order collocation algorithm. The collocation system is solved using a damped Newton method with an affine-invariant criterion function, as described in [18]. The equations for each cortex are solved in parallel, taking the current position of neighbouring cortices as possible binding locations. This process is repeated, updating the rest length of the cortices at every time step, until a global equilibrium is reached. Adaptive mesh refinement was used at every relaxation step, to ensure that node spacing remained numerically stable and fast. Additional nodes were added in regions where the mesh spacing was greater than 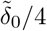 and nodes were removed where the spacing was less than 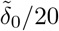. The model parameters used for the simulations are given in Table 1.

Source code for running the published simulations can be found at github.com/Alexander-Nestor-Bergmann/appcom. Documentation and quick-start tutorials can be be found at appcom.readthedocs.io.

